# Cell type-specific master metabolic regulators of Alzheimer’s disease

**DOI:** 10.1101/2025.07.11.664443

**Authors:** Yunguang Qiu, Yuan Hou, Liam Wetzel, Jessica Z.K. Caldwell, Xiongwei Zhu, Andrew A. Pieper, Tian Liu, Feixiong Cheng

## Abstract

Alzheimer’s disease (AD) exhibits metabolic heterogeneity; yet, the consequences on metabolic dynamics in a cell-type-specific manner and the underlying metabolite-sensor network basis remain unclear. Here, we show that neurons exhibit a striking decrease in energy and lipid-related metabolic activity, contrasted by an increase in microglial metabolism associated with neuroinflammation. To identify cell-type specific master metabolic regulators of AD underlying the metabolic alterations in AD, we introduce scFUMES (**<u>s</u>**ingle **<u>c</u>**ell **<u>FU</u>**nctional **<u>ME</u>**tabolite-**<u>S</u>**ensor), an algorithm integrating single-cell RNA sequencing, interactomics, genomics, transcriptomics, and metabolomics from human brain biobanks. Applied to two AD-vulnerable regions (middle temporal gyrus and dorsolateral prefrontal cortex), scFUMES uncovers hundreds of AD-associated regulators, with neurons and microglia showing the most interactions. Particularly, scFUMES pinpoints genetics-informed master metabolic regulators across AD severity, sex and *APOE* genotype (e.g., PPARD-glycerol in microglia). Experimental testing reveals that two interaction pairs predicted by scFUMES, neuronal palmitic acid bound fatty acid binding protein 3 and gut metabolite indole-3-propionic acid binding to kynurenine aminotransferase 1, both lower pathological tau species in AD. Collectively, scFUMES systematically maps AD master metabolic regulators, offering insights into cellular metabolic heterogeneity and therapeutic strategies for AD and other AD-related dementia if broadly applied.

## INTRODUCTION

Alzheimer’s disease (AD) is a multifactorial neurodegenerative disorder marked by heterogeneous dysregulation of cellular metabolism^1, 2^. While prior metabolomics and genomics studies have linked metabolites to genetic regulators^3^, mechanistic insights into cell-type-specific metabolic reprogramming (e.g., metabolite-sensor network) underlying AD pathogenesis and disease progression remain limited. To date, these include amyloid-β induced metabolic shifts in microglia from oxidative phosphorylation to glycolysis^4^ and AD-associated astrocytic profiles^5^. Metabolic activity alterations rewire metabolite-sensor network in cell metabolism. For example, a master metabolic regulator, glucose interacts with GLUT1 in astrocytes while with GLUT3 in neurons^6^. Increasing glucose uptake in neurons, but not astrocytes, has been suggested to alleviate AD pathology in *Drosophila*^7^. Despite these, a comprehensive understanding of metabolite-sensor networks across diverse cell states in the AD brain is lacking.

Metabolites serve dual roles as substrates in biochemical pathways and as signaling molecules regulating cellular processes through interactions with enzymes and receptors^8^. For example, gut-derived metabolites have emerged as influential modulators of disease pathways across multiple disorders^9, 10^. Advances in metabolomics^11^ and *in vivo* functional assays^12^ have begun to map the transient, low-affinity interactions between metabolites and their sensors, including G-protein-coupled receptors (GPCRs)^13, 14^. However, the dynamic nature of these interactions in the context of diseases poses significant challenges for systematic characterization.

To address this gap, we developed **<u>s</u>**ingle **<u>c</u>**ell **<u>FU</u>**nctional **<u>ME</u>**tabolite-**<u>S</u>**ensor (scFUMES), a computational framework integrating single-cell/nucleus RNA sequencing (sc/sn-RNA seq) and functional interactome, combined with multi-omics data. Sc/sn-RNA seq combined with systems biology analysis have opened new possibilities to characterize cell-type-specific heterogeneity in health and AD^15–17^. Our scFUMES identified biologically cell-type-specific master metabolic regulators and quantified metabolic activity alterations between neurons and glial cells in AD (e.g., energy and lipid metabolism). Specifically applied to two AD-vulnerable brain regions (middle temporal gyrus (MTG) and dorsolateral prefrontal cortex (DLPFC)), scFUMES prioritized genetics-informed metabolite-sensor networks tied to AD severity, sex differences, and *APOE* genotype combined with Mendelian randomization (MR). Neurons and microglia exhibited the most pronounced alterations, with microglia displaying particularly high metabolic diversity (e.g., PPARD-glycerol). Experimental validation confirmed that two scFUMES-predicted master metabolic regulator pairs (fatty acid binding protein 3 (FABP3)-palmitic acid and kynurenine aminotransferase 1 (KYAT1)-indole-3-propionic acid) both reduced the accumulation of pathological tau species in AD.

## RESULTS

### A systems biology framework to quantify metabolic heterogeneity in AD

To mechanistically unravel cell-type-specific metabolic heterogeneity at a transcriptome-wide scale, we established a systems biology analysis of single-cell transcriptomics, interactomics, genomics, and metabolomics from AD brains with deep phenotypic profiles (**Figure 1**). In this framework, we evaluated the metabolic activity alterations in AD by quantifying the metabolic disorders and pathway changes based on metabolic gene expression profiles from single-nucleus RNA-seq (snRNA-seq) (**Figure 1a and b**). Alterations of metabolic genes in AD impact the signaling metabolites behaviors^3^. We posit that AD pathologies disrupt the brain metabolic activities (e.g., bioenergetic metabolism^18^) and thereby rewire the metabolite-sensor network. Thus, we next presented scFUMES, a computational algorithm to characterize cell-type-specific metabolite-sensor interactions in AD **(Figure 1c**). The levels of certain metabolites are linked to genetic predisposition to AD^19^. We then prioritized AD genetics-informed master metabolic regulators by integrating scFUMES with Mendelian Randomization (MR) analysis (**Figure 1d**) and investigated their paired sensors across AD phenotypes and risk factors, including disease severity, sex, and *APOE* status (**Figure 1e**). Finally, we experimentally validated the functional roles of scFUMES-predicted metabolite-sensor pairs using tau cellular models (**Figure 1e**).

**Figure 1.**
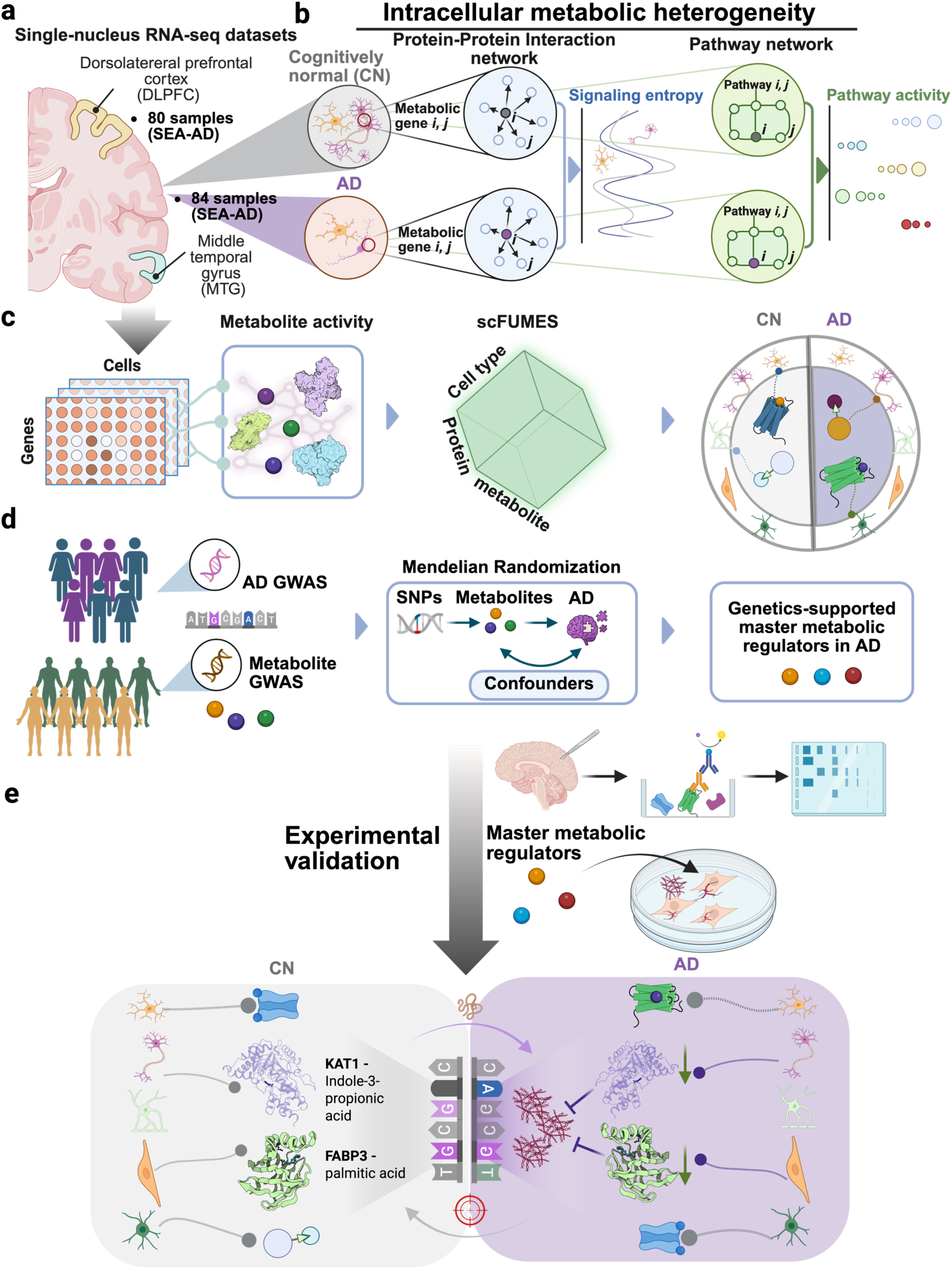
A systematic framework for the characterization of metabolic heterogeneity in AD. **a** Single sc/sn RNA-seq datasets. Datasets were retrieved from The Seattle Alzheimer’s Disease Brain Cell Atlas (SEA-AD), including ∼1.2 million nucleus from dorsolateral prefrontal cortex (DLPFC) region in 80 participants and ∼1.3 million nucleus from middle temporal gyrus (MTG) region in 84 participants. **b** Schematic diagrams showing intracellular metabolic heterogeneity calculation. The computation of single-cell signaling entropy and pathway activity based on the gene expression profiles in either protein-protein interaction (PPI) networks or metabolic pathway networks. The significant differences were scored for AD compared to non-AD. **c** Schematic of **scFUMES** analysis. Circular metabolites were collected from HMDB databases followed by filtering and cleaning. Metabolites functional/physical activities were comprehensively retrieved from ChEMBL and BindingDB databases. Single-cell gene expression matrix and metabolite-sensor activities were utilized to predict and score metabolite-sensor pairs in a given cell type in different disease pathologies. **d** Schematic of Mendelian Randomization (MR) genetics analysis. Instrumental variables (IVs) for each metabolite were selected from recently large-scale metabolite GWAS datasets. Four AD GWAS summary statistic datasets were treated as the outcome. Four multiple SNPs-based MR methods were adopted to estimate causal effects of circular metabolites on AD. **e** Integrating analysis of genetics and single cell-specific metabolite-sensor pairs. Combined the results from scFUMES (**d**) and MR analysis (**e**) to infer and validate genetics-supported single-cell-specific metabolite-sensor network across AD phenotypes, including brain regions, AD severity, sex difference, and APOE4 phenotypes.

### Preferentially elevated metabolic heterogeneity in AD microglia but not neurons

To assess the cellular metabolic heterogeneity associated with AD neuropathologic change [ADNC]), we firstly quantify the cell-type-specific metabolic activity by inferred a metabolic signaling entropy^17^. Signaling entropy calculations were based on relative transcriptomic gene expressions in the context of interaction network, and high signaling entropy indicates high metabolic activity^17, 20^. We estimated the metabolic activities for ∼1.3 million snRNA-seq profiles from the MTG brain region and ∼1.2 million snRNA-seq profiles from the DLPFC from The Seattle Alzheimer’s Disease Brain Cell Atlas (SEA-AD, *cf.* **Methods**)^21^. We chose these regions because of the MTG’s known AD vulnerability^22^ and the DLPFC’s role in impaired synaptic plasticity in AD^23^. In the MTG, neuronal cells (including excitatory neurons (ExN) and inhibitory neurons (InN)) had the highest metabolic activity, but it dropped the most in AD compared to non-AD individuals (**Figure 2a**). Glial cells showed elevated metabolic activity in AD than non-AD except for astrocytes (Ast). For example, microglia attained the lowest metabolic activity in non-AD, however, which significantly increased in AD (**Figure 2a**). Consistent trends were also observed in the DLPFC (**Figure 2b**), except for oligodendrocytes (ODC), indicating region-specific dysregulations and a role of ODC in early AD^24^.

**Figure 2.**
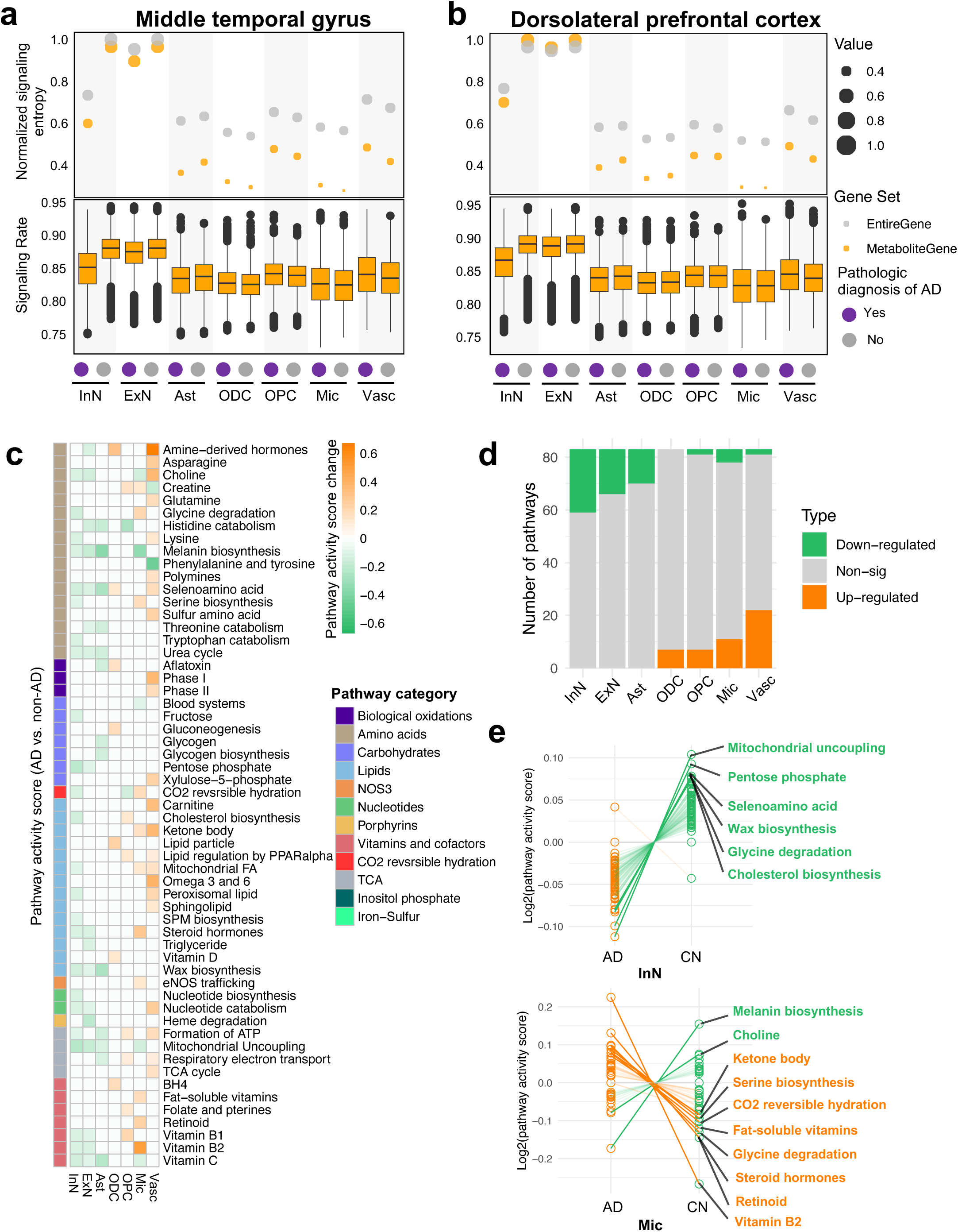
Molecular profiles of cell-type-specific metabolic heterogeneity in AD. **a,b** Single-cell-specific signaling entropy for AD and non-AD samples in the middle temporal gyrus (MTG, **a**) and dorsolateral prefrontal cortex (DLPFC, **b**) brain regions. The lower sub-panels showing metabolic signaling rates (SR), representing the cellular entropy rates over the weighted metabolic PPI networks within specific cell types. Elevated signaling rates indicate high uncertainty or potential disorders. The upper dot plots showing normalized signaling entropy based on signaling rate values, where orange indicates metabolic signaling entropy (considering only metabolic related genes) and gray represents signaling entropy for all human genes. **c** Pathway activity scores for metabolic pathways across cell types in the MTG region. Differences between AD and non-AD that are greater than 0.1 indicate significant changes. Orange indicates up-regulated pathways in AD, while green indicates the pathways that are down-regulated in AD. **d** Summary of the number of significant metabolic pathways in AD as illustrated in **c**. **e** Exemplar metabolic pathways in InN and microglia.

We next turned to inspect the metabolic heterogeneity using a pathway activity scores (AlzPAS) based on weighted gene expression alterations in 83 metabolic pathways (**Figure 2c-e and S1, Table S2**)^25, 26^. Compared to non-AD, both MTG and DLPFC exhibited distinct metabolic pathway profiles across multiple cell types in AD (**Figure 2c and S1a**). For example, glycogen biosynthesis was found to be significantly and specifically down-regulated in AD astrocytes, consistent with decreased glycolysis^27^ and glycogen metabolism^28^ in AD astrocytes. We next compared the pathway distributions (**Figure 2d and S1b**). Consistent with above metabolic activity analysis (**Figure 2, a and b**), metabolic pathways enriched in neurons and astrocytes were significantly downregulated in AD (*e.g.,* inhibitory neurons, 24/24, *p* < 0.01) in both brain regions. By contrast, microglia exhibited more upregulated pathways in microglia (11/16). Specifically, we observed downregulated mitochondrial uncoupling and cholesterol biosynthesis pathways specifically in neurons (**Figure 2e and S1c**), two key pathways associated with bioenergetic metabolism^29^ or APOE-mediated lipid regulation^30^ in AD. In microglia, however, we found that neuroinflammatory-regulated pathways, such as ketone body^31^ and Vitamin B2^32^, were up-regulated (**Figure 2e and S1c**).

Compared to the MTG, we found amino acid metabolism (asparagine and threonine catabolism) were specifically changed in inhibitory neurons from DLPFC, revealing brain region-specificity. Likewise, triglyceride metabolism is specifically upregulated in the DLPFC microglia, a potential pathway contributing to microglial activation and the subsequent pro-inflammation^33^. In summary, these observations reveal cell type-specific metabolic heterogeneity in AD, exemplified by neurons and microglia.

### Discovery of neuron/microglia-specific metabolite-sensor networks in AD

To mechanistically understand the cellular metabolic heterogeneity in AD, we next sought to investigate metabolite-sensor pairs in a cell type-specific manner. We sourced 2,647 experimentally determined metabolite-sensor pairs connecting 509 signaling metabolites and 517 proteins (**Figure S2a**, *cf.* **Methods**). We excluded natural products, drugs, and exogenous metabolites to focus on cellular metabolism. Organic acid (31%, 159/509) and lipids (29%, 148/509) were the most abundant metabolites (**Figure S2b**), while enzymes (34%, 176/517) and GPCRs (17%, 87/517) were the most common sensors (**Figure S2c**). We developed an algorithm, termed **<u>S</u>**ingle **<u>C</u>**ell **<u>Fu</u>**nctionally **<u>ME</u>**tabolite-**<u>S</u>**ensor communication (scFUMES), to prioritize biologically relevant metabolite–sensor pairs from snRNA-seq data (*cf.* **Methods** and **Figure 1c**). Via scFUMES, we inferred the interaction strength per metabolite-pair with cell type-specificity through cell label shuffling (termed alzMET score). As shown in **Figure 3**, 410 cell type-specific pairs are distinct in AD in the MTG region (alzMET score difference > 0.5 between AD and non-AD, FDR < 0.05, **Table S3**). Of these, neurons and microglia exhibited the highest number of distinct metabolite-sensor pairs, sensors, and metabolites (**Figure S2d**), aligning with our metabolic activity analysis (**Figure 2**).

**Figure 3.**
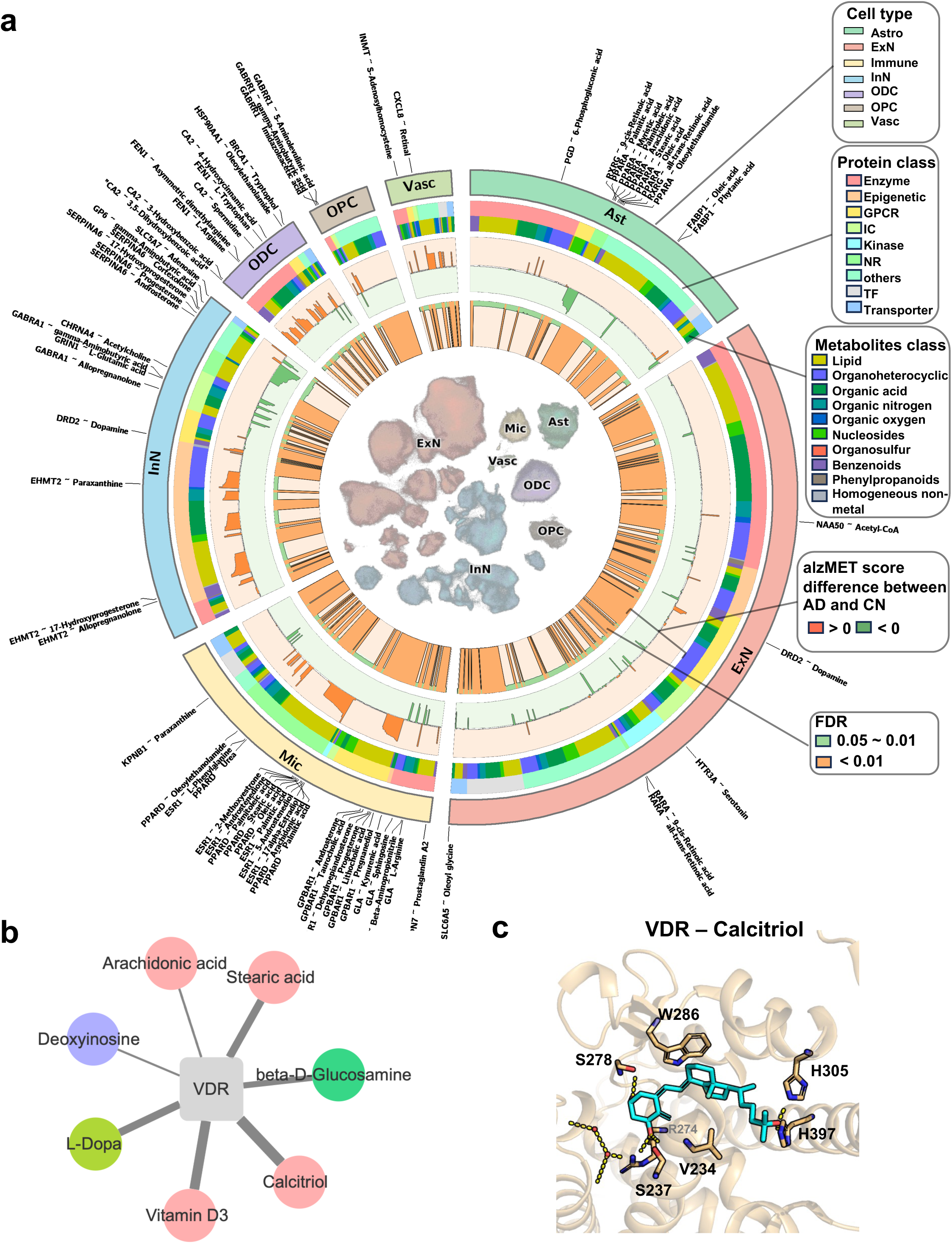
Systematic characterization of metabolite-sensor network in AD. **a** Circos plot showing overall metabolite-sensor pairs identified by scFUMES for each cell-type in AD versus non-AD samples in the MTG. UMAP for single-cell distributions was shown in the center of panel. The innermost circle represents False discovery rate (FDR) values, where orange indicates FDR < 0.01, green indicate 0.01 < FDR < 0.05. The second and third circles indicate communication score differences between AD and non-AD as calculated by scFUMES algorithm. Green showed pairs specific to non-AD, while orange indicates AD-specific pairs. The fourth circle categorizes the metabolites, and the fifth circle categorizes protein targets. The outermost circle showed the cell types corresponding to the UMAP. Labels highlight the highly selective, cell-type-specific pairs (|communication score difference| > 10). Phenotype-specific metabolite-sensor pairs are defined as those with |communication score difference| > 0.5. **b** Network depicting VDR-associated metabolite-sensor pairs in microglia. **c** 3D complex structure of VDR paired with Calcitriol (PDB ID: 7QPP).

We next investigated the cell-specific metabolic sensors involved in biological metabolite-sensor pairs. Totally, 97 cell-type specific sensors (metabolite-regulated proteins) associated with AD were prioritized (**Figure S2, e and f**). 45.4% (44/97) of the sensors have previously been reported to be associated with AD (**Table S3**). Particularly, 28% (27/97) AD-associated sensors were involved in metabolic pathways (**Figure 2c**), suggesting crucial roles of metabolites on mediating signaling pathways^8^. 12 sensors were specific to microglia, such as AD-associated peroxisome proliferator-activated receptor delta (PPARD) and vitamin D receptor (VDR) (**Figure S2, e and f**). Specifically, we found that VDR interacts with seven metabolites, including vitamin D3 and its active metabolite calcitriol (**Figure 3b**), with 3D crystal structure^34^ showing that calcitriol forms strong hydrophobic interactions with VDR (K_d_ = 0.065 nM). Vitamin D pathway was specifically upregulated in AD microglia (**Figure S1**). Consistently, VDR was more highly expressed in microglia in AD (log_2_FC = 0.80, **Figure S3a**). These data indicate specific activation of microglial VDR in AD^35^.

Using scFUMES, we also revealed that free fatty acid receptor 3 (FFAR3)-short-chain fatty acids (SCFAs) are highly specific in oligodendrocytes cells (FFAR3 paired with acetic acid, butyric acid, and propionic acid, **Figure S2g**). FFAR3 mediates biological effects of SCFAs produced by gut microbiota^36^, and FFAR3 deficiency has been suggested to exacerbate neuroinflammation and early aging in mouse brain^37^. Butyric acid was significantly prioritized to interact with FFAR3 in non-AD groups (alzMET score = -0.74, FDR < 0.01) and potently activated FFAR3 via a strong electrostatic interaction (**Figure S2h**, EC_50_ = 12 μM). Consistent with these results, FFAR3 expression was reduced in oligodendrocytes in AD human brain (log_2_FC = - 0.56, **Figure S3b**). Collectively, we prioritized cell-specific variations of biological metabolite-sensor pairs by scFUMES.

### Long-chain fatty acid palmitic acid reduces Tau neuropathology via FABP3

Among scFUMES-predicted sensors associated with AD, Fatty Acid Binding Protein 3 (FABP3, a key binding enzyme for palmitic acid) was strongly associated with AD in excitatory neurons (alzMET score = 0.72, FDR of 0.03, **Figure S2, e and f**). We also inspected *FABP3* gene expression alterations across different brain regions and cell types (**Figure S3, c and d**). The RNA level of *FABP3* was significant reduced in excitatory neurons (AD vs. non-AD, log_2_FC = -0.11, FDR = 1.58 x 10^-44^, **Figure S3c**) and throughout multiple brain regions (e.g., superior temporal gyrus, **Figure S3d**). This is consistent with the previous studies that reduced level of FABP3 in the brain may contribute to AD-like pathology^38^.

Protein level changes of FABP3 in the human cerebrospinal fluid and brain have been implicated in tau pathologies, and genetic elimination of *FABP3* has been shown to contribute to AD in mice^38, 39^. Therefore, we next experimentally inspected the roles of scFUMES-predicted palmitic acid or FABP3 in Tau pathology. The protein level of FABP3 was found to be significantly decreased in AD brains (*p* = 0.047, **Figure 4, b and c, Table S4**) compared to cognitive health controls using Western blot and immunohistochemistry staining analysis of 20 postmortem brains (*cf*. **Methods**). We further investigated the effects of FABP3 and its signaling metabolite palmitic acid on cellular Tau seeding activity, a process involving the recruitment and misfolding of normal tau molecules by tau aggregates in tauopathies^40^. We found that the overexpression of FABP3 in TauRD P301S FRET Biosensor (TauRD) cells, following treatment with tau seeds extracted from the brains of P301S-Tau mice, markedly reduced Tau seeding activity (*p* = 1.1 x 10^-5^, **Figure 4d**). Our results further showed that increasing FABP3 significantly decreased RIPA-insoluble hyperphosphorylated-tau 231 (pTau-231) but not RIPA-soluble pTau-231 (*p* = 5 x 10^-4^ for insoluble pTau-231, *p* = 0.056 for soluble pTau-231, **Figure S4**).

**Figure 4.**
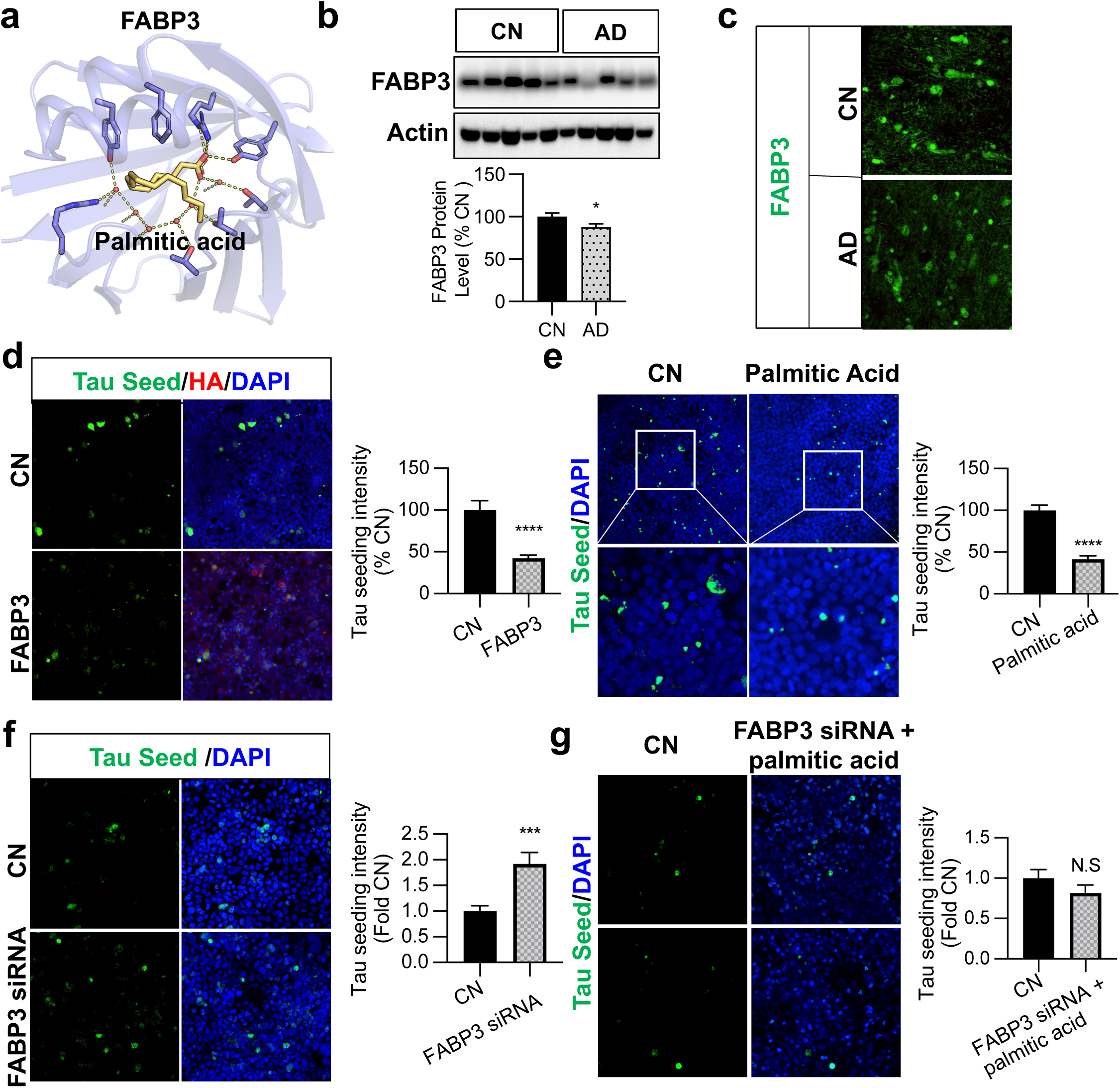
scFUMES-predicted palmitic acid reduces Tau neuropathology via FABP3. **a** 3D Binding mode of FABP3-palmitic acid. **b** Representative blots of RIPA-soluble lysates of cognitively normal (CN) and AD cortical brains subjected to immunoblotting for FABP3, and actin (versus CN, *p* = 0.047, n = 10 CN, n = 10 AD). **c** Representative images of postmortem frontal cortex brain sections from CN and AD cases immunostained for FABP3 (green) (versus CN, *p* = 0.0091, n = 5 CN, n = 5 AD). **d** TauRD cells transfected with vector control and FABP3-HA with the treatment tau seed for 24h and subjected to ICC for HA (FABP3-HA) and direct fluorescence for intracellular tau seed. Tau seed (green) intensity was quantified as tau seeding index (*p* =1.1 x 10^-5^, n = 4 replicates). **e** TauRD cells pretreated with 200mM of palmitic acid for 12h, with 36h Tau seed treatment, were subjected to ICC for tau seeding activity. (*p* < 0.0001, n = 3 replicates). **f** TauRD cells transfected with control and FABP3 siRNA, with the treatment of Tau seed for 24h subjected to ICC for endogenous FABP3 (red) and direct fluorescence tau seed (green). Tau seed (green) intensity was quantified as tau seeding index (*p* = 4 x 10^-4^, n = 4 replicates). **g** TauRD cells transfected with control and FABP3 siRNA, with the pretreatment of 200mM of palmitic acid for 12h and later treatment of Tau seed for 24h subjected to ICC for Tau seed activity. Tau seed (green) intensity was quantified as tau seeding index (*p* = 0.22, n = 3 replicates). Unpaired t test was utilized to determine significance. *, *p* < 0.05; **, *p* < 0.01; ***, *p* < 0.001; ****, *p* < 0.0001; ns, not significant.

Lastly, we examined whether palmitic acid plays a protective role by virtue of its interaction with FABP3. Palmitic acid, a long-chain fatty acid, binds to FABP3 with a K_d_ of 1μM^41^ (**Figure 4a**), and was found to significantly decrease the Tau seeding activity (*p* < 1 x 10^-5^, **Figure 4e**). Here, knockdown of FABP3 using FABP3 siRNA significantly increased the Tau seed activity (*p* = 4 x 10^-4^, **Figure 4f**) and abolished the protective effect of palmitic acid (*p* = 0.22, **Figure 4g**), consistent with our model. In summary, we demonstrated that a scFUMES-predicted pair involving FABP3 and its signaling metabolite palmitic acid both alleviate tau pathology *in vitro*.

### Discover likely causal master metabolic regulators in AD

We next investigated genetics-supported master metabolic regulators in AD using MR approaches (cf. **Methods**). Four recent metabolite genome-wide association studies (GWAS)^42–45^ and three AD GWAS datasets^46–48^ were utilized to estimate the causal effects of metabolites in AD. In total, 124 metabolites were identified with likely causal effects on AD risk (FDR < 0.05, **Figure S5, Table S5**). Of these, 62.1% (77/124) were prioritized in the largest AD GWAS dataset^46^ (**Figure 5a**). Lipids (n = 53, 42.7%) and amino acids (n = 36, 29.0%) were the top two categories (**Figure 5b**). Specifically, directionality analysis showed that ∼60% of metabolites (71/124) were associated with reduced risk of AD (**Figure 5b**), such as epiandrosterone sulfate (an androgen metabolite, **Figure 5c**), whereas 44 out of 124 metabolites (e.g., glucose, tyrosine and triglyceride) associated with elevated risks of AD (**Figure 5c**).

**Figure 5.**
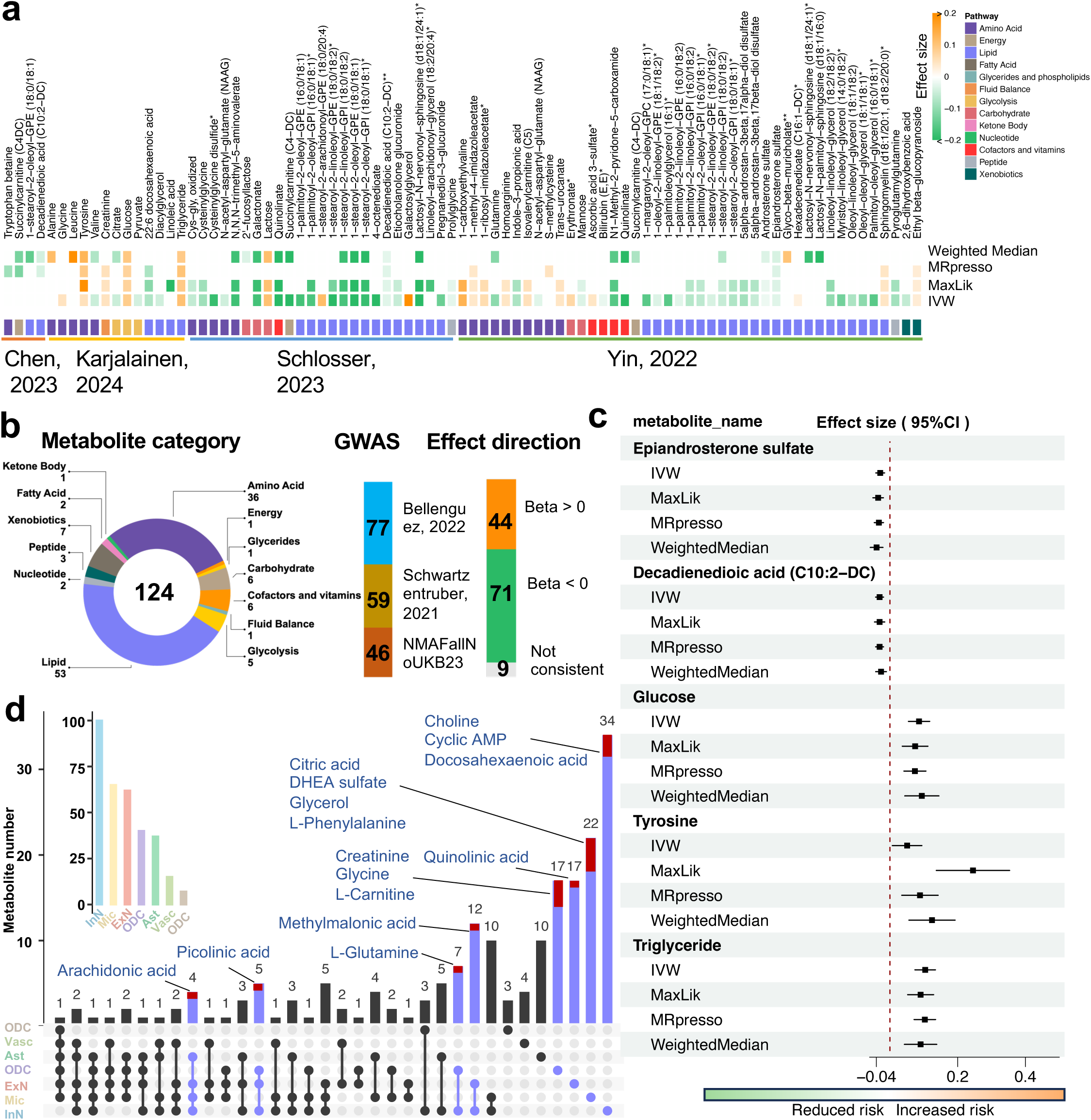
Discovery of AD likely causal master metabolic regulators across cell types. **a** Prediction of AD likely causal metabolite by utilizing Mendelian Randomization (MR) analysis. Four latest metabolite GWAS and 1 AD GWAS datasets were employed. Four MR methods were adopted, including IVW, MaxLik, MRpresso and Weighted Median. Significance, FDR < 0.05. **b** Summary of the number of metabolites prioritized from MR analysis based on their categories, GWAS studies and causal effects. **c** Selected AD likely causal metabolites prioritized by MR analysis. The causal effects are consistent across four MR methods. The positive values indicate the increased risks of AD, while negative values indicate reduced risks of AD. **d** Matrix for all intersections of cell types sorted by metabolite numbers. The overlapped AD likely casual metabolites are highlighted in red and labeled.

Next, we intersected the AD likely causal metabolites with the scFUMES-predicted networks. In the MTG, inhibitory neurons has the largest number of AD-associated metabolites (100), followed by microglia (64) (**Figure 3a and 5d**). Of these, 34 were specific to inhibitory neurons and 22 to microglia. Via chemical identity mapping, 15 AD likely causal metabolites were prioritized in the scFUMES-predicted metabolite-sensor network (**Figure 5d**). For example, a higher level of microglial phenylalanine is associated with increased risk of AD (β = 0.83). Across cell types, 26.6% (4/15) of likely causal metabolites were shared in various cell types. For example, arachidonic acid, a mediator of inflammation that contributes to Aβ production and AD pathogenesis^49^, was predicted to associate with increased risk of AD in both neurons and microglia (**Figure 5d**).

### A genetically informed metabolite-sensor network specific to neurons/microglia in AD

We next examined scFUMES-predicted metabolite-sensor pairs possessing AD likely causal metabolites (**Figure 3a and 5d**). In the MTG, scFUMES prioritized 27 such pairs, including 16 sensors, whereas 23 pairs (11 sensors) were identified in the DLPFC (**Figure 6, a and b, Table S6**). Of these, 9 sensors were specific to the MTG and 5 to the DLPFC, such as serine racemase (SRR), an enzyme in serine biosynthesis (**Figure 6a**) known to be elevated in the APP/PS1 AD mouse model^50^. SRR is also specifically paired with glycine in MTG oligodendrocytes, a metabolic regulator associated with increased risk of AD (**Figure 5a**). Conversely, we found that the DLPFC-specific farnesoid X receptor (FXR, encoded by the *NR1H4* gene, **Figure 6b**), which regulates bile synthesis and Aβ-related pathologies^51^, paired with an AD likely protective metabolite citric acid (**Figure 5a**) in InN cells. Furthermore, 4 pairs were conserved in both MTG and DLPFC with AD associations (**Figure 6, a and b**), including CHRNA4-choline (InN), VDR-arachidonic acid (microglia), GNAS-methylmalonic acid (InN) and TSG101-methylmalonic acid (ExN).

**Figure 6.**
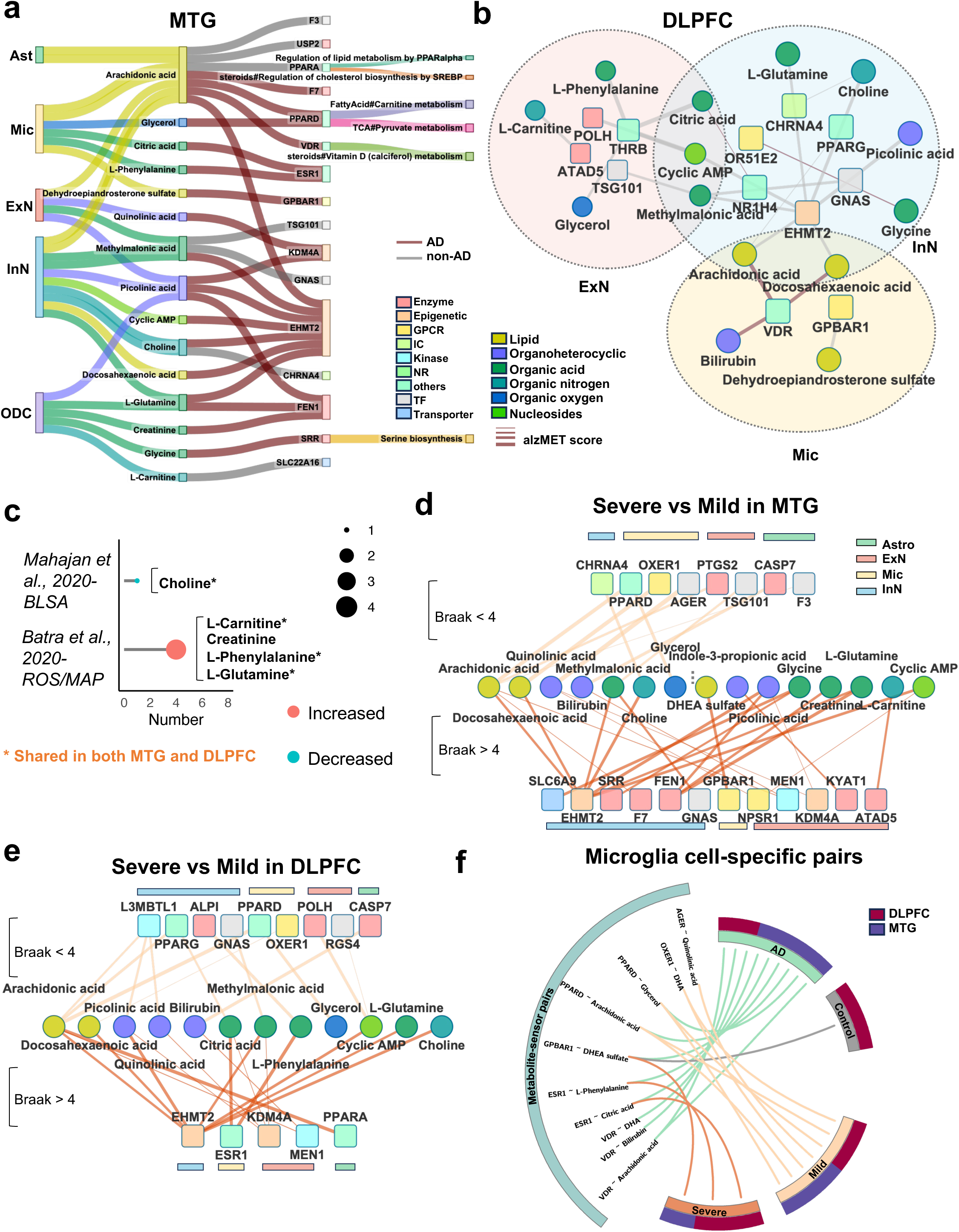
AD genetically informed metabolites-mediated metabolite-sensor network. **a,b** Network representations of genetics-supported metabolite-sensor pairs across cell types in MTG region (**a**) and DLPFC region (**b**). The colors for squares or circles denote cell types, metabolite and protein categories. The colors for lines denote AD or non-AD. **c** dot plots showing the significant altered metabolites in AD retrieved from two previous metabolomics studies (BLSA study^73^ and ROSMAP study^1^, |communication score difference| > 0.5, FDR < 0.05). **d,e** Network representations of genetics-supported metabolite-sensor pairs compared severe stage of AD to mild stage of AD in MTG region (**d**) and DLPFC region (**e**). **f** Circos plot showing microglia-specific metabolite-sensor pairs across groups (AD and non-AD, Severe AD and mild AD, MTG and DLPFC).

To evaluate whether the genetically supported master metabolic regulators were perturbed in AD cohorts, we re-analyzed two large-scale metabolomics datasets from the Baltimore Longitudinal Study of Aging (BLSA)^52^ and the Religious Order Study and Rush Memory and Aging Project (ROS/MAP)^1^. We found that 5 likely causal metabolites prioritized by high alzMET scores, with significantly altered levels in AD (FDR < 0.05), including choline, L-carnitine, creatinine, L-phenylalanine and L-glutamine (**Figure 6c, Table S7**). Among these, choline has been previously reported as significantly decreased in AD^52^, and we found it to be significantly downregulated in neurons and microglia across both the MTG and DLPFC (**Figure 2c and S1a**). Our MR and scFUMES analysis indicated that choline may have AD protective effects by interacting with acetylcholine receptor subunit alpha-4 (encoded by *CHRNA4*) in InN cells (β = - 0.50, Ki = 7 μM, **Figure 6, a and b**)^53^.

As metabolic dysregulations have been implicated in AD progression^54^, we next investigated the differences between severe (Braak stage > 4) and mild (Braak stage < 4) AD pathology. In the MTG, severe AD displayed greater disruption than mild AD (23 versus 9 pairs, 23 versus 9, **Figure 6d).** Astrocyte-restricted metabolite-sensor pairs were found exclusively in mild AD, indicating early metabolic dysregulation in astrocytes. By contrast, severe AD was dominated by epigenetic sensor (*e.g.*, KDM4A and EHMT2)-mediated metabolite-sensor pairs (40%, 9/23), indicating a key role of epigenetic regulation in severe AD. Specifically, the gut microbiota-derived metabolic regulator indole-3-propionic acid (IPA) was linked to AD severity (**Figure 6d**). Notably, our scFUMES and MR analysis suggested that a higher level of IPA is associated with reduced AD risk by engaging kynurenine aminotransferase 1 (KYAT1) in ExN (**Figures, 3a and 5a**). Consistent with metabolomics analysis, significantly increased metabolites in AD, including L-carnitine, creatinine, and L-glutamine, are associated with severe AD (**Figure 6c**). In the DLPFC, the number of AD severity-related metabolite-sensor pairs is comparable (13 and 16, respectively, **Figure 6e**). 16 metabolite-sensor pairs were shared between MTG and DLPFC, such as EHMT2-choline. Despite this, two metabolites (citric acid and L-phenylalanine) were specific to the DLPFC.

Specifically, we systematically analyzed the genetics-supported metabolite-sensor pairs in microglia. 10 unique pairs were significantly specific to microglia, consisting of 6 sensors and 8 metabolites (**Figure 6f**). The MTG region revealed twice as many microglial metabolite-sensor pairs as the DLPFC in AD. As shown in **Figure 3b**, VDR is a key sensor in microglia. Our MR and scFUMES suggested that inflammation-associated arachidonic acid^55^ may be associated with a high risk of AD by activating VDR (AC_50_ = 6.78 μM) in microglia. The endogenous VDR activator vitamin D could increase Aβ deposition and exacerbate AD^56^. In parallel, L-phenylalanine is a master metabolic regulator in microglia and our analysis of metabolomics data showed that L-phenylalanine is significantly increased in AD (**Figure 6c**), consistent with our MR analysis (β = 0.8) and previous studies^57^.

Given that sex and *APOE4* are two strong risk factors for late-onset AD^58^, we next examined differences across sexes and *APOE* genotypes. We observed that females exhibited more genetically supported metabolite-sensor pairs than males (12 vs. 7) across most cell types in the MTG (**Figure S9, a and b, Table S8**), while the opposite trend was observed in the DLPFC (4 vs. 13; **Figure S10a, Table S9**). Of these, two are specific to female microglia, including PPARD-arachidonic acid and PPARD-glycerol (**Figure S9c**), which are also associated with AD severity (**Figure 6f**). Our MR analysis suggested that high levels of arachidonic acid and glycerol are associated with high AD risks (**Figure 5a**).

*APOE4* carriers exhibited a similar number of genetically supported metabolite-sensor pairs, with non-*APOE4* carriers in both MTG (13 versus 14 pairs) and DLPFC (12 versus 18 pairs) (**Figures S9d, S10b and S11b**). 69% of *APOE4*-specific pairs (9/13) were conserved across brain regions. Specifically, in the MTG, *APOE4-*specific pairs clustered in InN, while non-*APOE4-*specific pairs distributed to other cell types (**Figure S9e, Table S8 and S9**). Notably, two pairs specific to microglia in non-APOE4 carriers (PPARD-arachidonic acid and PPARD-glycerol) were also female-specific (**Figure S9f**). These results suggest that sex differences and APOE4 phenotypes shape distinct metabolite-sensor profiles in neurons and microglia.

### Gut metabolite indole-3-propionic acid reduces Tau phosphorylation via KYAT1

We next sought to functionally interrogate the scFUMES-predicted, AD severity-related, genetics-supported metabolite-sensor pair, indole-3-propionic acid (IPA)-KYAT1 (**Figure 6d**). A previous clinical study indicated that gut microbiota-produced IPA mediated neuroprotective effects in elderly^59^. Additionally, KYAT1 is a key enzyme in the production of kynurenic acid^60^, high levels of which are associated with slowed AD progression^61^. To test whether IPA alleviated AD pathology through regulating KYAT1, we first established a direct strong interaction between them (**Figure 7a**, PDB ID: 3FVU^62^). Experimentally, we found that IPA significantly reduced the Tau seeding activity (*p* < 1 x 10^-4^, **Figure 7b, Table S10).** We also assessed the protein changes of KYAT1 by western blot and immunohistochemistry, which revealed significantly decreased KYAT1 in AD brains (*p* = 4.2 x 10^-3^, **Figure 7, c and d**). Importantly, this aligned with bulk-transcriptomic data analysis showing significantly decreased KYAT1 in AD prefrontal cortex of males and posterior cingulate cortex in both sexes (**Figure S8, Table S11**). Importantly, increasing KYAT1 significantly reduced the Tau seeding activity (*p* = 1 x 10^-6^, **Figure 7e**), as well as insoluble pTau-231 (*p* = 0.041, **Figure S4**). Next, we inspected the effects of IPA on AD pathologies via KYAT1. Knockdown of KYAT1 in vitro significantly reduced KYAT1 protein (*p* < 1 x 10^-4^, **Figure S7**) and accelerated Tau seeding activity (*p* = 5 x 10^-6^, **Figure 7f**). Furthermore, knockdown of KYAT1 also significantly impeded the protective effect of IPA (**Figure 7, b and g,** *p* = 0.036), indicating that the gut metabolite IPA may alleviate AD Tau pathologies by sensing KYAT1.

**Figure 7.**
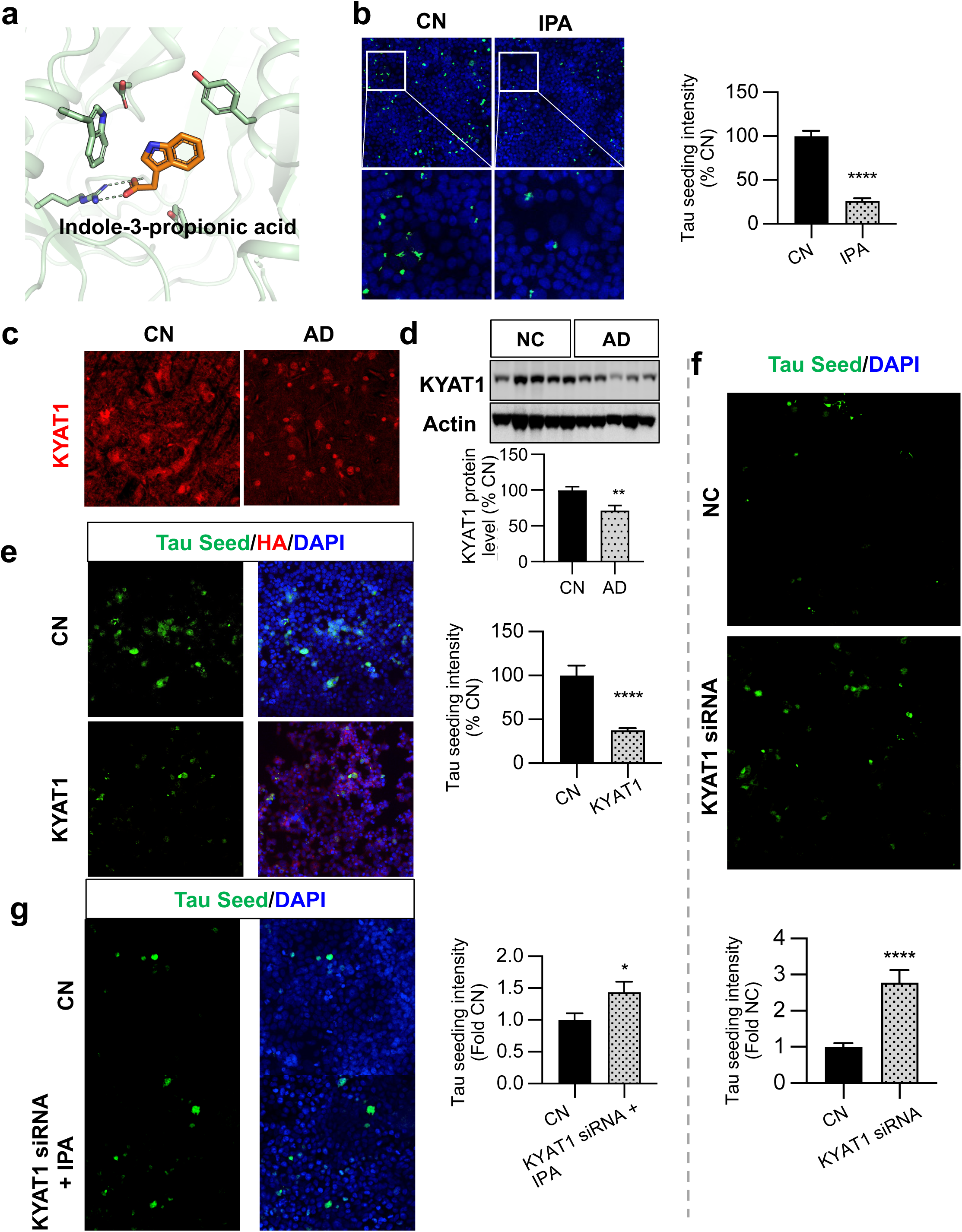
Gut metabolite indole-3-propionic acid reduce Tau phosphorylation via KYAT1. **a** 3D Binding mode of KYAT-indole-3-propionic acid (IPA). **b** TauRD cells pretreated with 200mM of indole-3-propionic acid for 12h with 36h Tau seed treatment, were subjected to ICC for tau seeding activity. (*p* < 0.0001, n = 3 replicates). **c** Representative images of postmortem frontal cortex brain sections from CN and AD cases immunostained for KYAT1 (red) and DAPI (blue) (versus CN, *p* = 0.0083, n = 5 CN, n = 5 AD). **d** Representative blots of RIPA-soluble lysates of CN and AD cortical brains subjected to immunoblotting for KYAT1, and actin (versus CN, *p* = 0.0042, n = 10 CN, n = 10 AD). **e** TauRD cells were transfected with vector control and KYAT1-HA, with the treatment of tau seed for 24h, and subjected to ICC for HA (KYAT1-HA) and direct fluorescence for intracellular tau seed. Tau seed (green) intensity was quantified as tau seeding index (*p* = 1 x 10^-6^, n = replicates). **f** TauRD cells transfected with control and KYAT1 siRNA, with the treatment of Tau seed for 24h subjected to ICC for endogenous KYAT1 (red) and direct fluorescence tau seed (green). Tau seed (green) intensity was quantified as tau seeding index (*p* = 5 x 10^-6^, n = 4 replicates). **g** TauRD cells transfected with control and KYAT1 siRNA, with the pretreatment of 200mM of indole-3-propionic acid for 12h and later treatment of Tau seed for 24h subjected to ICC for Tau seed activity. Tau seed (green) intensity was quantified as tau seeding index (*p* = 0.036, n = 3 replicates). Unpaired t test was utilized to determine significance. *, *p* < 0.05; **, *p* < 0.01; ***, *p* < 0.001; ****, *p* < 0.0001; ns, not significant.

In summary, our integrated genetics and bioactivity-based master metabolic regulators analysis highlights dynamic cellular maps of metabolite-sensor pairs across neurons and microglial in different brain regions with respect to AD severity and risk factors. Our findings may provide clues to explain intricate metabolic heterogeneity in AD in ways that lead to the discovery of new metabolism-based therapeutics for AD.

## DISCUSSION

AD is marked by profound metabolic dysregulation^1,63^, yet the mechanisms linking metabolite-sensor interactions to cellular pathology remain poorly understood. Our study addresses this gap through a systematic framework that integrates a statistical algorithm scFUMES combined with metabolomics, and genetic data. By investigating two AD-vulnerable brain regions, we uncovered metabolic activity changes in cell type-specific manner. Neurons exhibited depletion of metabolic pathways (energy or lipid metabolism)^16^, while microglia displayed heightened metabolic plasticity, likely reflecting their inflammatory adaptation to disease states^64^. These findings align with established AD hallmarks of neurodegeneration and neuroinflammation, while advancing a metabolic perspective on their interplay.

scFUMES uniquely quantifies metabolite-sensor network perturbations, revealing genetic and phenotypic drivers of AD progression. For example, choline-CHRNA4 interactions linked to AD-associated SNPs underscore genetically influenced metabolic disruptions. The framework further disentangled risk factor-specific heterogeneity, including APOE4-driven microglial PPARD-glycerol interactions and sex-dependent metabolic reprogramming. Based on the alzMET score, we highlighted several potential AD-related master metabolic regulators, such as palmitic acid interacting with FABP3. FABP3 has been associated with Tau pathologies^65^. In addition, DRD2 was also prioritized in neurons associated with AD, an AD GABAergic neuron marker^66^ paired with dopamine. Another AD-associated sensor, euchromatic histone lysine methyltransferase 2 (EHMT2), possesses the most significant metabolite-sensor pairs in inhibitory neurons, such as S-Adenosylhomocysteine-EHMT2 (K_i_ = 570 nM). High level of EHMT2 has been detected in AD, and the inhibition of EHMT2 may rescue the cognitive functions in AD^67^. Glucose is engaged in glycolysis pathway, and a high level of blood glucose is associated with increased risks of AD, consistent with a previous finding^68^.

Gut microbial metabolites hold potential for AD^10^. We clarified a potential AD protective pair in vitro that includes gut microbiota-derived master metabolic regulator indole-3-propionic acid pairing with KYAT1. Apart from it, we also observed several pairs interacting with SCFAs that are associated with non-AD. For example, FFAR3-acetic acid, FFAR3-butyric acid, HDAC8-butyric acid, and FFAR3-propionic acid, suggesting their potential protective effects in AD^69^. Additionally, the complex roles of SCFAs in modulating AD pathology may be attributed to their cell type-specific effects^69^. From the target perspective, GPCR was showed to be one of the largest potential AD targets following the enzyme^70^ (**Figure S2c**). For example, we found that an immune-related chemokine receptor 6 (CCR6) was significantly prioritized to interact with sphingosine in ExN in AD (IC_50_ = 20.9 μM, Figure 3a). CCR6 has been a potential target for treating chronic inflammatory diseases^71^ and a biomarker for AD in a mouse model^72^.

While scFUMES provides a foundation for mapping cell type-specific metabolic networks, its predictive power is currently constrained by the scope of existing data sets, particularly the sparse availability of metabolite-protein interaction data and region-specific metabolite concentrations. Furthermore, future *in vivo* validation of prioritized pairs, alongside integration of metabolic flux models, will refine the translations utility of this new approach.

In conclusion, scFUMES pioneers a systems-level strategy to decode metabolic heterogeneity in AD, providing a potential bridge between molecular mechanisms and new metabolite-targeted therapy. As multi-omics datasets expand, this framework will accelerate the identification of master metabolic regulators across various diseases and conditions, offering direction for precision therapy targeting metabolic dysfunction for preserving and restoring brain health.

## ACKNOWLEDGMENTS

This work was primarily supported by the National Institute of Aging (NIA) under Award Number R01AG084250, R56AG074001, U01AG073323, R01AG066707, R01AG076448, R01AG082118, RF1AG082211, and R21AG083003, and the National Institute of Neurological Disorders and Stroke (NINDS) under Award Number RF1NS133812 to F.C. AAP was supported by The Valour Foundation and Department of Veterans Affairs Merit Award I01BX005976. AAP is also supported as the Rebecca E. Barchas, MD, Professor in Translational Psychiatry of Case Western Reserve University and the Morley-Mather Chair in Neuropsychiatry of University Hospitals of Cleveland Medical Center. AAP also acknowledges support from NIH/NIA RO1AGs066707, NIH/NIA 1 U01 AG073323, the Louis Stokes VA Medical Center resources and facilities. This work was supported by grants from the National Institutes of Health (NIH) [R01AG086365 to T.L, R03AG084948 to T.L]. T.L was also supported by the Research Education component of the Cleveland Alzheimer’s Disease Research Center (NIA P30 AG072959).

## AUTHOR CONTRIBUTIONS

F.C., T.L., and Y.Q. conceived the study. Y. Q. performed computational analysis. L.W. performed in vitro experiments. Y.Q., Y.H. and L.W. performed all data analysis. X.Z., J.Z.K.C., A.A.P. discussed and interpreted the results. Y.Q. and T.L. wrote the manuscript. F.C., Y.Q., and T.L. revised the manuscript. All authors have read and approved the final manuscript.

## METHODS

### SEA-AD single-nucleus cohort

Large-scale single-cell RNA-seq data were utilized from The Seattle Alzheimer’s Disease Brain Cell Atlas (SEA-AD) study cohort (Synapse ID: syn26223298), including two brain regions, middle temporal gyrus (MTG)^24^ and dorsolateral prefrontal cortex (DLPFC)^21^. A total of 1.38 million nuclei from post-mortem MTG brain region from 84 participants and 1.22 million nuclei from post-mortem DLPFC brain region from 80 participants were isolated. Two brain regions are originated from the 80 participants. Annotation of cell types (astrocytes, excitatory neurons, inhibitory neurons, microglia, oligodendrocytes, OPCs and Vascular cells) was referred to the previous study^21^. Gene expression counts in cells were harmonized and normalized and UMAP was generated by utilizing Scanpy package^74^. For the purpose of this study, details of AD pathological metadata for SEA-AD cohorts were summarized, including age at death, sex, APOE4 status, cognitive status, CERAD score, ADNC score, Braak stage, and Thal phase.

### Metabolic signaling entropy analysis

AD-associated cell type-specific Metabolic Signaling Entropy (alzMSR) algorithm is built based on a previous study^17^. 11,751 genes involved in protein-protein interactions were retrieved from Pathway Commons^75^. Specifically, 2,130 metabolic genes were collected from Reactome^76^ to build a metabolic protein-protein network. Through gene mapping, a 1,604 x 1,604 metabolic interaction matrix were generated based on the entire protein-protein interaction network. The gene expressions were calculated by normalizing the gene counts for each single-nuclei dataset. Local signaling entropy indicates the uncertainty at each gene within a network (intracellular heterogeneity), while the global signaling entropy (signaling entropy rate, SR) quantifies the overall uncertainty or disorder of the entire network by considering the long-term behavior of the network at single-cell level. Briefly, for each gene 𝑖, compute its local signaling entropy 𝑆_𝑖_ as:

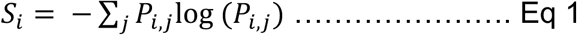

Where 𝑃_𝑖,*j*_ is the transition probability from gene 𝑖 to gene 𝑗. Next, a weighted sum of local signaling entropies is used to calculate an SR for each cell. A stationary distribution 𝑢_!_ was computed derived from gene’s expression matrix. Then SR was calculated:

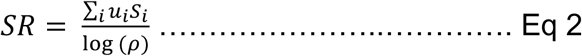

Where 𝜌 is largest eigenvalue of the adjacency matrix in the network. Through this calculation, the SR could be defined with the range of values between 0 – 1. The higher SR, the more metabolic heterogeneity of a specific cell has.

### Alzheimer’s metabolic pathway activity score

The AD metabolic pathway activity score^25^ was calculated based on previous studies^26^. Briefly, a total of 83 metabolic pathways were evaluated for normalized single-nuclei datasets across different brain regions. For each pathway, we weighted genes in the pathway based on its occurrence. For each comparation group, e.g., AD vs. non-AD, we calculated the relative gene expression for each gene per cell type by adjusting the global average gene expression. The Alzheimer’s pathway activity score (alzPAS) was computed as a weighted average of the adjusted AD-related relative gene expressions. Outliers in each pathway with relative expression levels that greater than three times 75th percentile or below 1/3 times 25th percentile were excluded. The pathway that has an alzPAS score > 1 was considered as enriched in a specific cell type. To achieve cell-type specific alzPAS score for each pathway, the cell labels were randomly shuffled 1000 times. *P* values were calculated derived from the permutation test and only significant deviations from the neutral baseline (score of 1) are retained for each pathway. The cutoff of alzPAS difference between groups (e.g., AD vs. non-AD) is 0.1.

### scFUMES algorithm

Cell-type specific Single Cell FUnctionally MEtabolite-Sensor communication (scFUMES) algorithm was constructed by leveraging the large-scale single-cell/nuclei sequencing data and knowledge-based metabolite-sensor pairs. 228,855 circular metabolites were retrieved from HMDB database^77^. Natural products, drug or drug metabolites, toxin pollutant and exogenous metabolites were excluded. Next, functional or physical binding activity measurements (K_d_, K_i_, IC_50_, EC_50_, AC_50_ and potency) were retrieved for the metabolites. Totally, 11,763 measurements from BindingDB^78^ and 55,444 measurements from ChEMBL^79^ were collected. Given the metabolic activity heterogeneity via diverse assays, stringent quality control was performed: (1) Activity outliers for each metabolite-protein pair with max/min value ratio that is more than 10 were removed. (2) Duplicates were removed. (3) PAINS compounds^80^ with alarm structures for metabolites were filtered. (3) IDs of metabolites standardized by chemical name, SMILES, INCHIKEY, HMDB ID and KEGG ID. UniProt IDs were used for protein IDs. (4) Gene category and metabolites sources were mapped. (5) Activity cutoff (bioactive activity < 1 mM) was employed. (6) manually inspected. Finally, 2,647 high-quality measurements were retrieved including 509 signaling metabolites and 517 protein targets.

Sc/nRNA-seq data was preprocessed, including removing cells with fewer than 200 genes expressed and genes expressed in fewer than 10 cells. Normalization and log transformation were performed on total counts. Only the genes that are involved in the metabolite-sensor interactions are considered. Per each group, e.g., AD and non-AD, for each cell type 𝑘 and gene 𝑔, average gene expression per cell type 𝜇 was calculated:

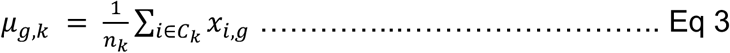

Where 𝑥_𝑖,*g*_ is the expression of gene g in cell 𝑖. Low expressed genes were filtered out (cutoff: 25^th^ percentile) to increase the statistical power. In addition, we also calculated the proportion of cells that each gene expressed. Next, we standardized the average gene expression across cell types to identify cell-type specific gene expression pattern:

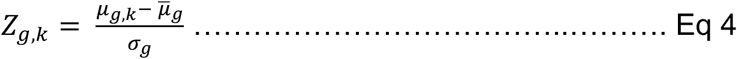

Where *μ̅*_*g*_. is the mean expression of gene 𝑔 per cell type, 𝜎_𝑔_ is standard deviation of gene 𝑔 per cell type.

For each metabolite-sensor pair (𝑔, 𝑚), we calculate a communication score for each cell type:

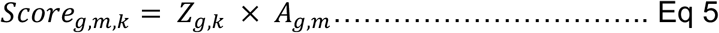

Where 𝐴*_g,m_* is the activity for each metabolite-sensor pair that indicate the potential interaction strength between metabolite and proteins in specific cell types. Here, the physical concentration of metabolites was assumed to be sufficiently high so that the concentration effects in the interactions could be neglected.

To assess the statistical significance of observed communication score, the cell labels were randomly shuffled, by default 1000 times to simulate a null distribution of communication scores. For each permutation, the score was recalculated, and *p* value was computed by the permutation test:

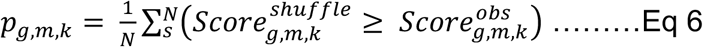

Where 𝑁 is the shuffle times. The *p* values were adjusted to control the false discovery rate (FDR) using the Benjamini-Hochberg procedure. The cutoff of significant score was FDR < 0.05. The significant differential pairs between AD and non-AD were determined if satisfying both FDR < 0.05 and |𝑆𝑐𝑜𝑟𝑒_AD_ − 𝑆𝑐𝑜𝑟𝑒_non-AD_| > 0.5.

### Mendelian randomization analysis

A summary data based Mendelian randomization (MR) analysis was performed to evaluate the potential causal effects of signaling metabolites for AD. Four recent metabolite genome-wide association studies (GWAS)^42–45^ and three AD GWAS datasets^46–48^ were retrieved from GWAS catalog (https://www.ebi.ac.uk/gwas/) or original papers. Totally, the GWAS for over 5,000 metabolites were included. Metabolite ratio or uncharacterized metabolites were removed. To select high-confidence and GWAS significant instrumental variables (IVs), two thresholds were applied: *p* < 5 x 10^-8^ and F statistics > 10. In addition, independent instruments were pruned by using PLINK (v1.9) clumping with r^2^ threshold of 0.1 and a specify an LD window of ± 500 kb. Variants were harmonized and two-sample MR was conducted by utilizing TwoSampleMR (https://mrcieu.github.io/TwoSampleMR/) under R (v4.3.0). To improve the statistical power, 4 MR models were employed that consider the number of proposed IVs ≥ 3: IVW, MR-PRESSO, MaxLik and Weighted Median^81^. For IVW and MaxLik, heterogeneity *p* value was calculated. FDR < 0.05 as the cutoff of effects of metabolite on AD. To improve the accuracy, we further pruned the AD likely causal metabolites by discarding metabolites with inconsistent effect directions in different MR methods.

### Gene expression analysis

For single-nucleus datasets, gene expression levels were calculated by adopting pseudobulk methods. The counts were aggregated for each cell type in a sample-level, producing pseudobulk samples that mirror bulk RNA-seq datasets. The gene expression was indicated by summation raw counts. For different groups, Log2Foldchange (log2FC) were calculated, e.g., AD and non-AD.

For bulk RNA-seq data, residualized gene counts was utilized for comparison. The data were retrieved from AD Knowledge Portal (syn21241740). The Datasets include source data derived from The Religious Order Study (ROS) and Memory and Aging Project (MAP), the Mayo Clinic Brain Bank and Mount Sinai/JJ Peters VA Medical Center NIH Brain and Tissue Repository (MSBB). To remove biological or technical covariates, such as batch effect or sample size, residualized gene counts were calculated by a linear model. Adjusted *p* value and log_2_FC were calculated. Adjusted p value < 0.05 was adopted as the statistically significant.

### Metabolomics data analysis

Two metabolomics studies that reported the metabolomics changes in BLSA study^73^ and ROSMAP study^1^ are collected, respectively. In the BLSA study, 30 samples from ITG brain region were used for metabolomics analysis. In the ROSMAP study, 514 samples from the DLPFC brain region were used for metabolomics. Only differential metabolites between AD and non-AD were analyzed based on the studies, FDR < 0.05.

### Structure analysis

3D structures of metabolite-sensor complex were retrieved from PDB databank (https://www.rcsb.org/). 3D cartoon diagrams were created by using PyMOL (v3.1.0).

### Human brain samples

Human cortical tissues utilized for immunoblotting experiments have been obtained from the Brain Bank at Case Western Reserve University and the NIH Neurobiobank. The neuropathology of normal controls (ages 60–92 years, PMI range 4–25 hours) and Alzheimer’s disease cases (ages 67–90 years, PMI range 3–26 hours) was assessed. All procedures followed to approved IRB protocols and complied with institutional bioethics guiding principles.

### Cell Culture

The HEK293 TauRD biosensor^82^ and Hela-V5-Tau stable cells^83^ were cultured in Dulbecco’s modified Eagle’s medium (DMEM, Thermo Scientific, MA, USA) supplemented with 10% fetal bovine serum (FBS) and 1% penicillin/streptomycin (P/S). The cells were incubated in an incubator at 37℃ in a humidified atmosphere with 5% CO2, as previously described^84^.

### Immunocytochemistry and Microscopy

Cells on the cover glass were washed with PBS and subsequently fixed with 4% paraformaldehyde (PFA) for 15 minutes at room temperature. Fixed cells were rinsed with PBS and then blocked with a blocking solution (0.2% Triton X-100, 3% normal goat serum) for 1 hour, followed by incubation with primary antibody at 4 °C overnight. The cells were then washed three times with PBS and incubated with Alexa-488 or Alexa-594 conjugated secondary IgG antibodies and DAPI for 1 hour at room temperature (Vector Laboratories, Burlingame, CA). The slides were subsequently washed three times with PBS and mounted using a fluorochrome mounting solution from Vector Laboratories. The images were captured with the Nikon AX Ti2 confocal microscope (Tokyo, Japan). All comparable images maintained equal laser power, exposure duration, and filter configurations. During picture acquisition and quantification, researchers were unaware of the experimental circumstances, and regions of interest were selected randomly. Adjustments to brightness and contrast were equally implemented across all comparable images.

### Immunoblotting

Brain tissues or HelaV5 cells were lysed using RIPA lysis buffer (50 mM Tris pH 7.4, 150 mM NaCl, 2 mM ethylenediaminetetraacetic acid (EDTA), 1% NP-40, 0.1% sodium dodecyl sulfate (SDS)). Total protein concentrations were quantified with a colorimetric detection kit (BCA Protein Assay, Pierce, USA). Equal amounts of protein lysates were assessed using sodium dodecyl sulfate–polyacrylamide gel electrophoresis and subsequently transferred to a nitrocellulose membrane (Millipore Corporation, Bedford, MA, USA). Targeted proteins were detected utilizing primary antibodies after a blocking step with 5% skim milk, followed by the application of peroxidase-conjugated secondary antibodies, and identified through ECL (Merck Millipore Corporation, Darmstadt, Germany). All immunoblot images were acquired using the LAS-4000 (GE Healthcare Biosciences, Pittsburgh, PA) and analyzed with ImageJ (NIH, Bethesda, MD).

### Antibodies and reagents

Anti-pTau231 (Catalog # 71429S) antibodies were procured from Cell Signaling (Danvers, MA, USA). β-actin (Catalog # 66009-1-Ig), anti-KYAT1 (Catalog# 30296-1-AP), anti-HA (Catalog# 81290-1-RR), and anti-FABP3 (Catalog# 60280-1-Ig) antibodies were procured from Proteintech (Rosemont, IL, USA). Palmitic acid (Catalog # 29558) was purchased from Cayman (Ann Arbor, MI, USA). Indole-3-propionic acid (Catalog # 57400-5G-F) was purchased from Millipore Sigma (Burlington, MA, USA).

### DNA plasmid, siRNAs, and Transfections

pPM-KYAT1-HA, pPM-FABP3-HA constructs were made by Genscript (Piscataway, NJ, USA). For DNA plasmid transfections, cells were transfected with Fugene HD (Promega, Madison, WI, USA) in Opti-MEM I (Invitrogen, Carlsbad, CA, USA) according to the manufacturer’s instructions and harvested 48h post-transfection. For siRNA transfections, Lipofectamine 2000 (Invitrogen, Carlsbad, CA, USA) was used according to the manufacturer’s instructions. The KYAT1 siRNA sequences are 5’-A.U.U.U.C.C.C.A.C.C.A.C.C.A.G.A.C.U.U.U.G.U.U -3’; and the FABP3 siRNA sequences are 5’ -A.C.A.C.U.U.G.U.G.C.G.G.G.A.G.C.U.A.A.U.U.U.U -3’. The siRNAs were purchased from Horizon Discovery (Waterbeach, UK).

## DATA AVAILABILITY

The source code for alzMSR is available via GitHub at: https://github.com/qiusir1/alzMSR. Other source codes will be available upon requests.

---

**Figure S1.**
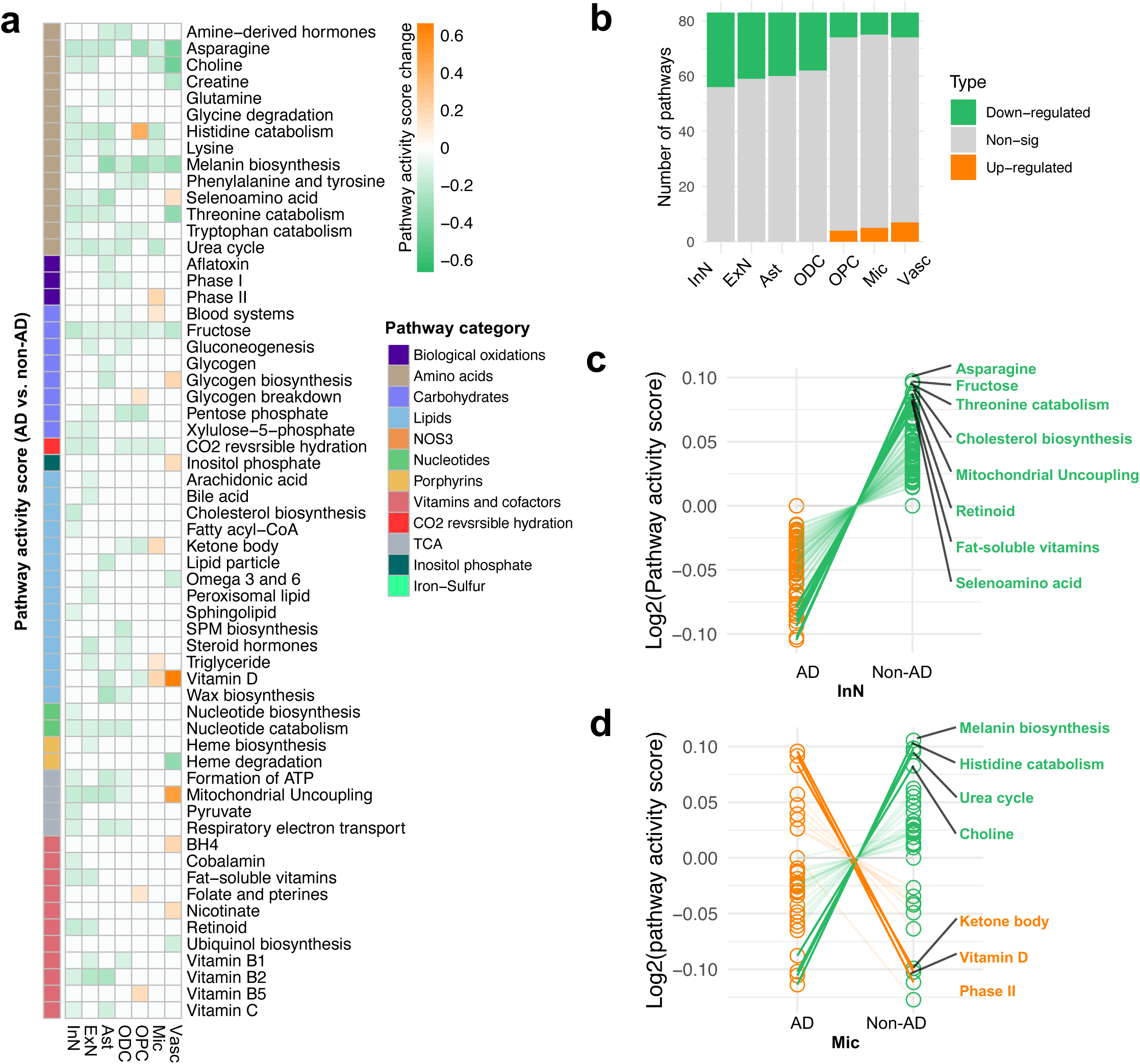
Profiling cell-specific metabolic heterogeneity in AD in the DLPFC. **a** Pathway activity scores for metabolic pathways across cell types in the DLPFC region. Differences between AD and non-AD that are greater than 0.1 indicates significant changes. Orange indicates up-regulated pathways in AD, while green indicates the pathways that are down-regulated in AD. **b** Summary of the number of significant metabolic pathways in AD as illustrated in **a**. **c** Exemplar metabolic pathways in InN and microglia. Figure colors and labels format are consistent with Figure 2.

**Figure S2.**
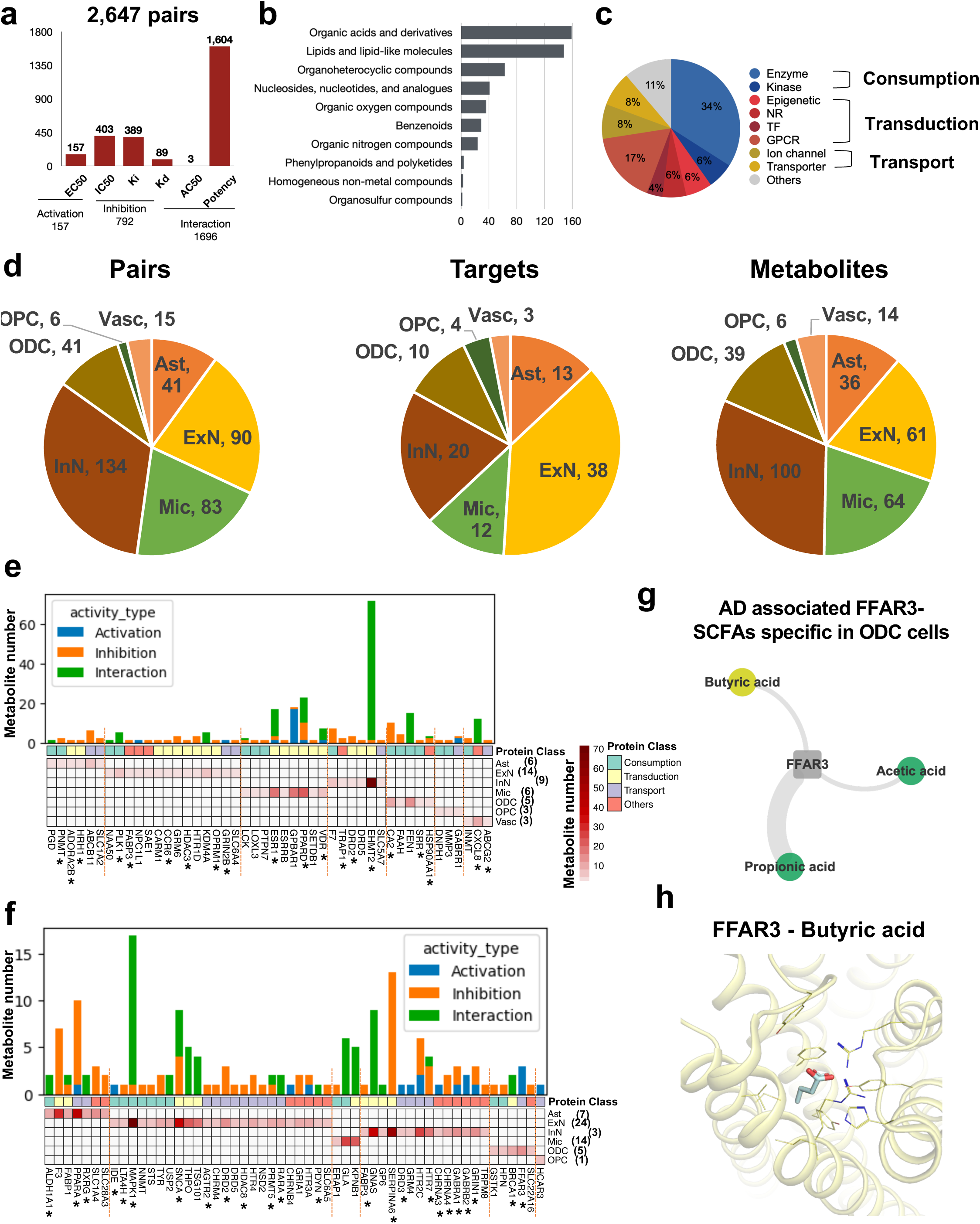
Summary of cell type-specific metabolite-sensor pairs from scFUMES. **a** Summary of metabolite-sensor physical/functional measurements retrieved from ChEMBL and BindingDB databases. **b,c** Categories of metabolites and targets from (a). **d** The numbers of pairs, targets and metabolites per cell types. **e,f** Target analysis across different cell types in AD (**e**) or non-AD (**f**). The targets that has been reported to associate with AD are denoted with “*”. The target numbers per each cell type are labeled in parentheses. The colors represent the number of metabolites for each target. **g,h** Exemplar 3D complex structures of FABP3-palmitic acid (**g,** PDB ID: 4TKJ) specific in AD ExN cells and FFAR3-butyric acid specific in non-AD ODC cells (**h,** PDB ID: 8J21).

**Figure S3.**
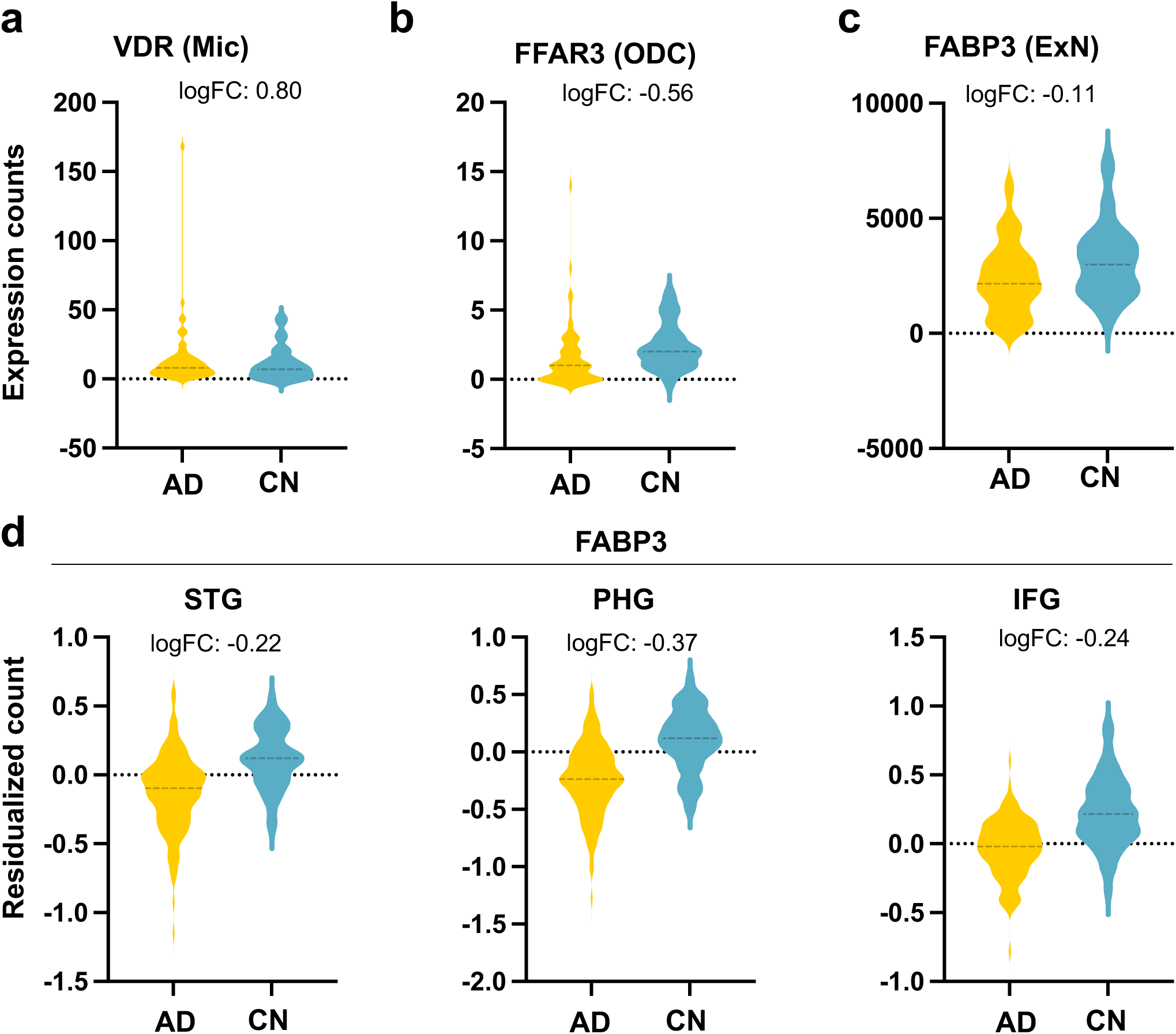
Gene expression alterations of selected proteins in different human brain regions and cell types. **a** FABP3 expression changes in brain regions (STG, PHG and IFG) at the transcriptomics RNA-seq level (AD versus non-AD). **b** Gene expression alterations of FABP3 in ExN, VDR in microglia and FFAR3 in ODC. Log2Foldchange (log_2_FC) were calculated compared AD to non-AD. Each point represents one individual.

**Figure S4.**
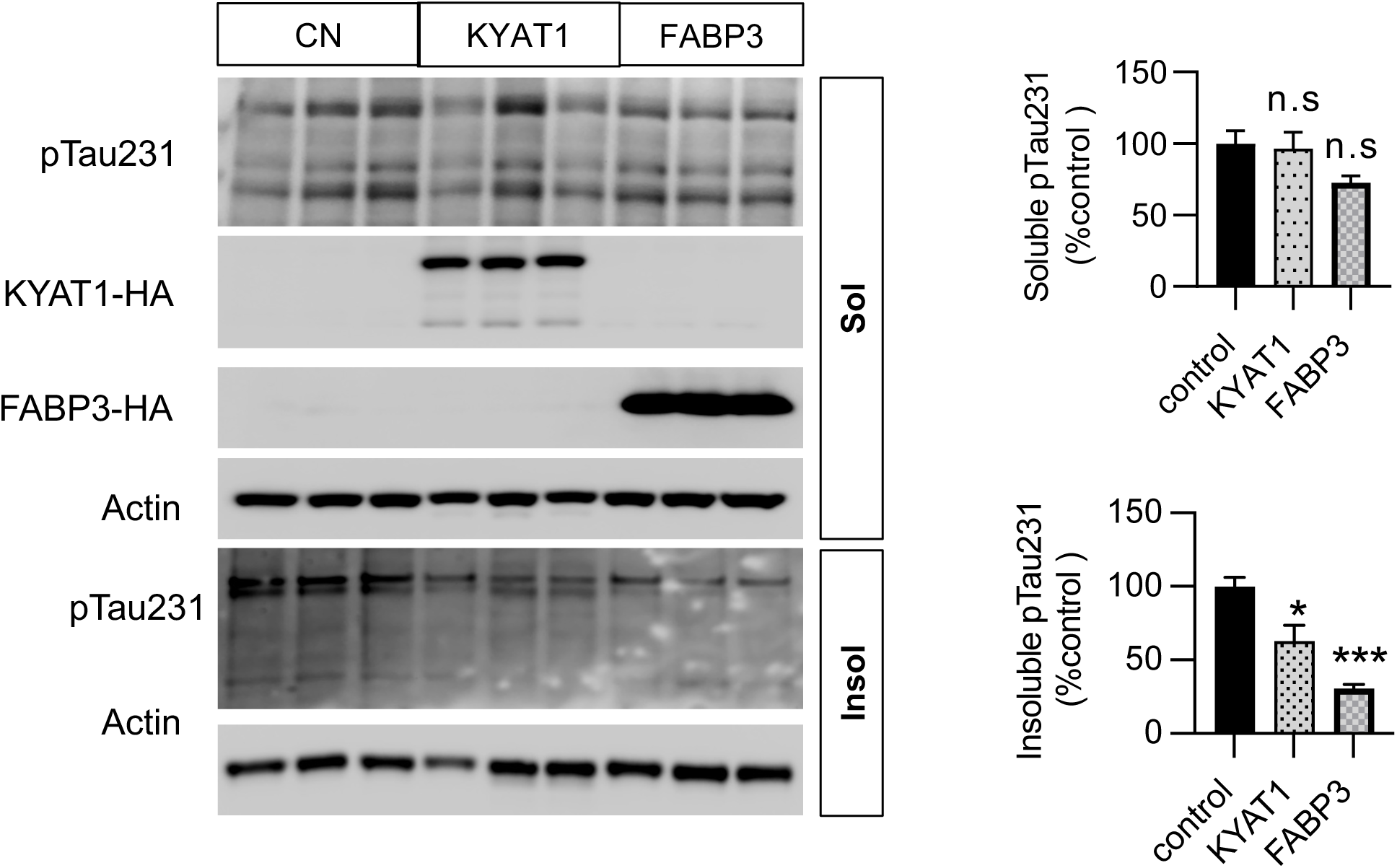
The functional effects of scFUMES-predicted FABP3 or KYAT1 on pTau-231. Western blot and quantitative analysis showing the overexpression of scFUMES-predicted FABP3 or KYAT1 significantly reduced insoluble pTau-231 (*p* = 5 x 10^-4^ for FABP3, *p* = 0.041 for KYAT1, n =3) while not soluble pTau-231 (*p* = 0.056 for FABP3, *p* = 0.82 for KYAT1, n =3 replicates). Unpaired t test was utilized to determine significance. *, *p* < 0.05; **, *p* < 0.01; ***, *p* < 0.001; ****, *p* < 0.0001; ns, not significant.

**Figure S5.**
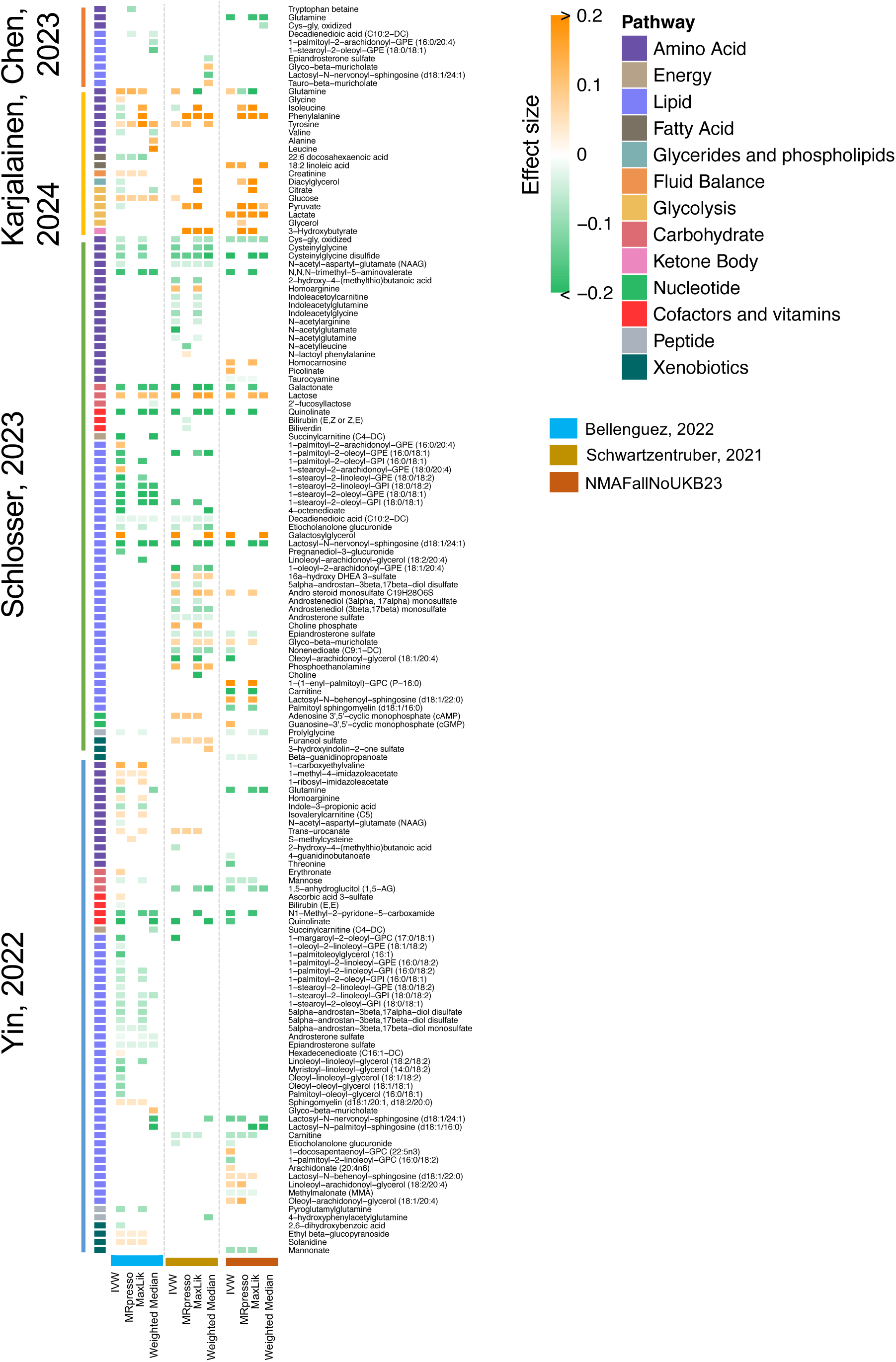
Summary of genetics-supported AD likely causal metabolites. Continue to Figure 4. We predicted AD likely causal metabolite based on four latest metabolite GWAS and three AD GWAS datasets were employed. Four MR methods were adopted, including IVW, MaxLik, MRpresso and Weighted Median. Significance, FDR < 0.05.

**Figure S6.**
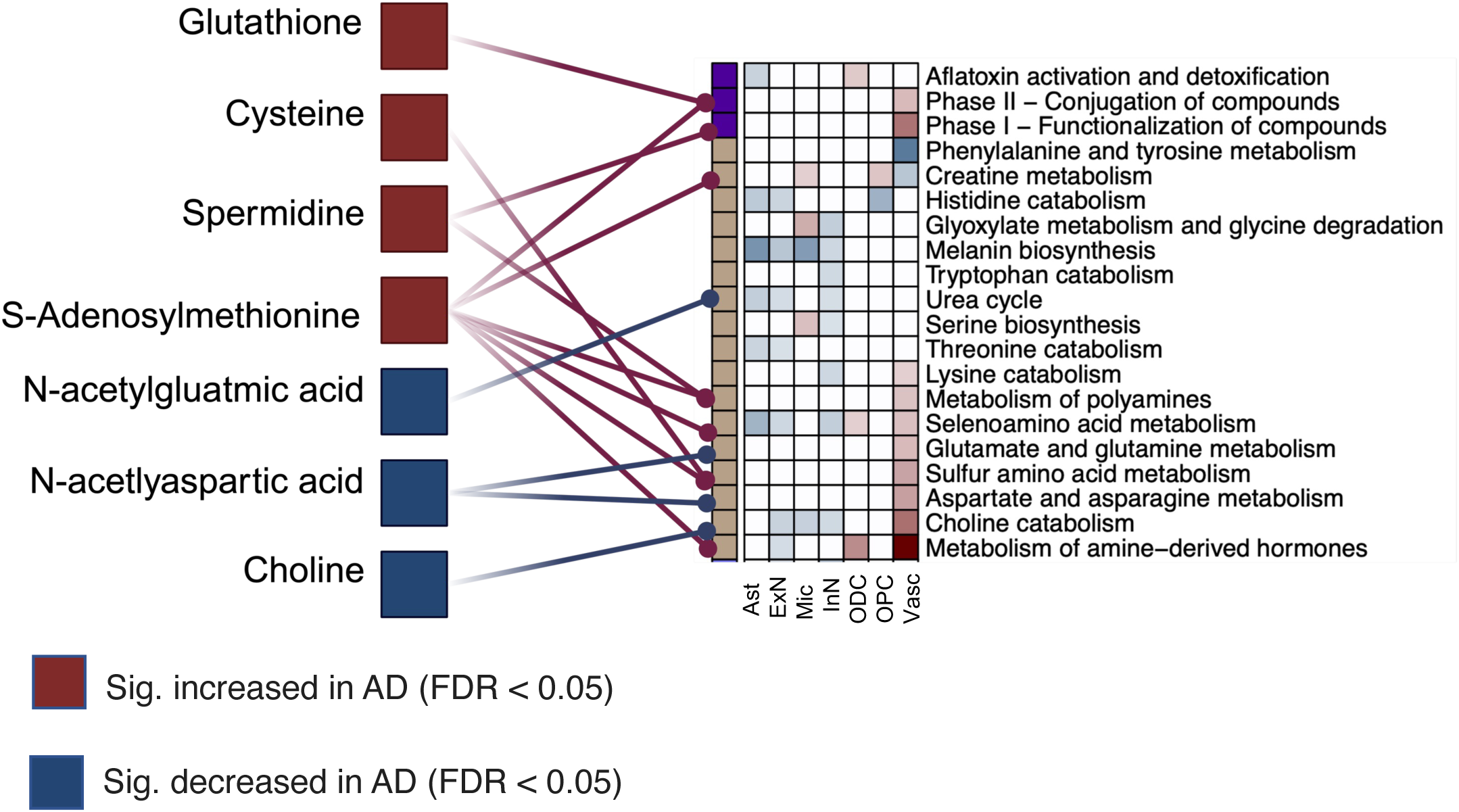
**Metabolic pathway analysis for AD-associated metabolites from the metabolomics studies**. Significant metabolites in AD were denoted in different colors (FDR < 0.05). Related metabolic pathways were connected to specific metabolites. Metabolic pathways were also seen in Figure 2c.

**Figure S7.**
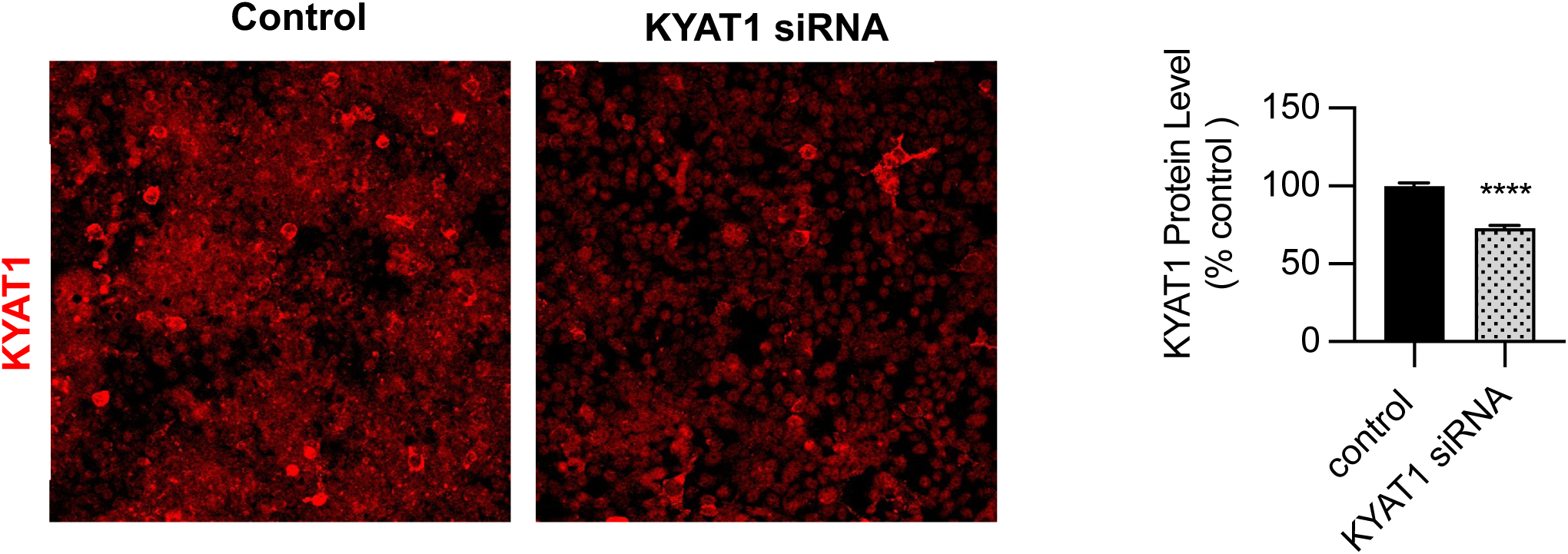
The protein level changes of KYAT1 knockdown in vitro. knockdown of KYAT1 via siRNA significantly reduce the KYAT1 protein level (p < 10^-4^). Unpaired t test was utilized to determine significance. *, *p* < 0.05; **, *p* < 0.01; ***, *p* < 0.001; ****, *p* < 0.0001; ns, not significant.

**Figure S8.**
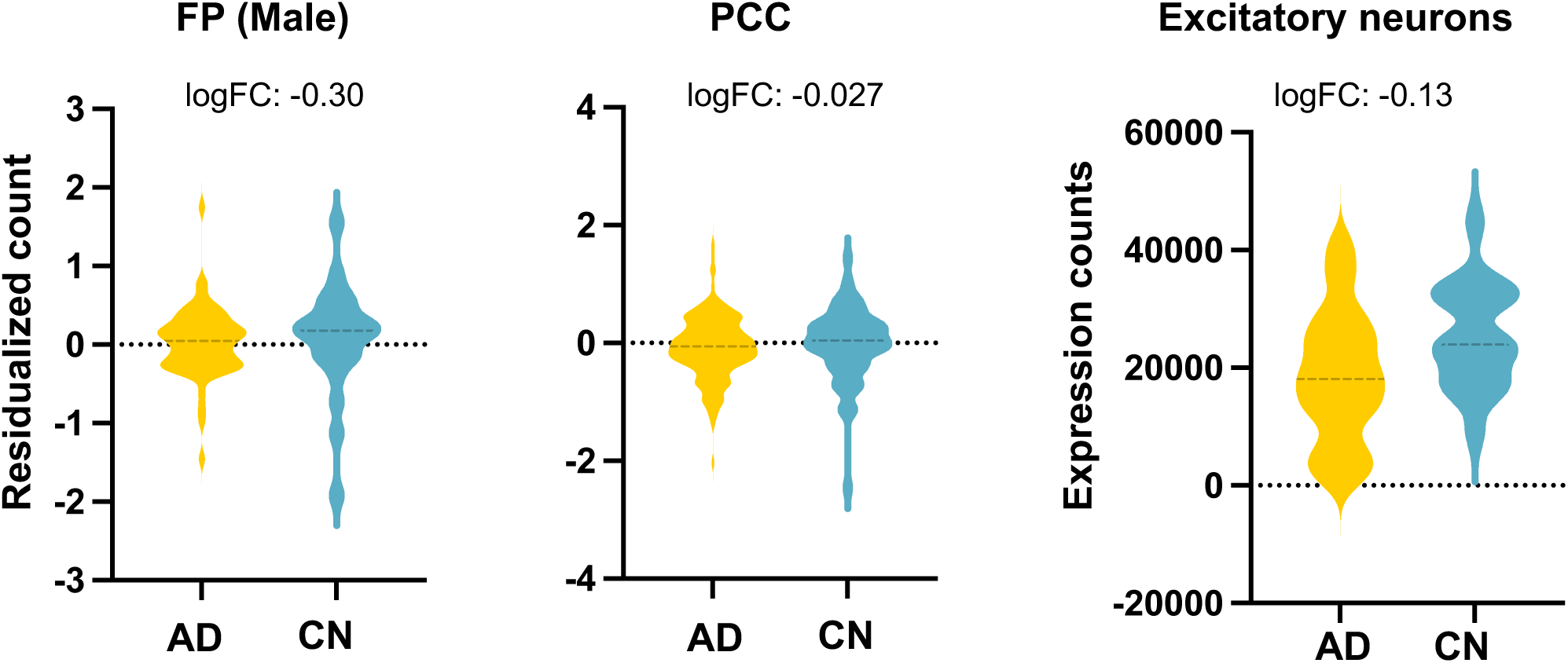
Gene expression alterations of KYAT1 in single cell or transcriptomic level. Differential expression of KYAT1 in FP (in male), PCC brain regions at bulk RNA-seq level and in ExN at single-cell level.

**Figure S9.**
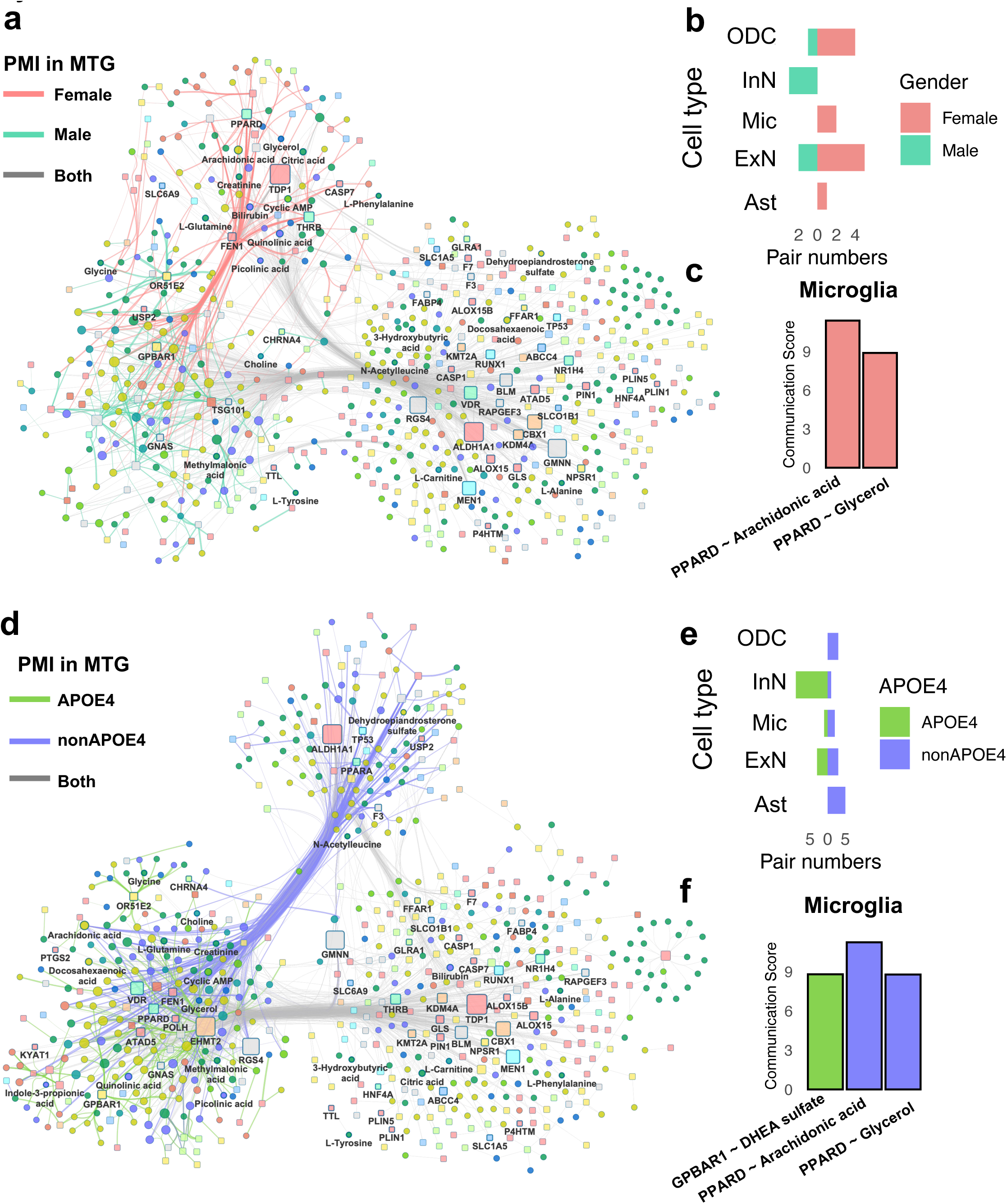
Genetic-supported metabolites-mediated metabolite-sensor network by AD risk factors in the MTG. **a** Network depicting sex-specific metabolite-sensor pairs. Line colors indicate the female or male-specific pairs. Metabolites are shown in circle, and proteins are shown in square. MR-prioritized metabolites and their pairs were labeled. **b** Histogram showing number of cell-specific genetic-supported metabolite-sensor pairs within female or male samples. **C** Microglia specific genetic-supported metabolite-sensor pairs. **d** Network depicting metabolite-sensor pairs in samples with *APOE4* or non-*APOE4* phenotypes. Line colors indicate the *APOE4* or non-*APOE4*-specific pairs. **e** Histogram showing number of cell-specific genetic-supported metabolite-sensor pairs within *APOE4* or non-*APOE4* samples. **f** Microglia specific genetic-supported metabolite-sensor pairs.

**Figure S10.**
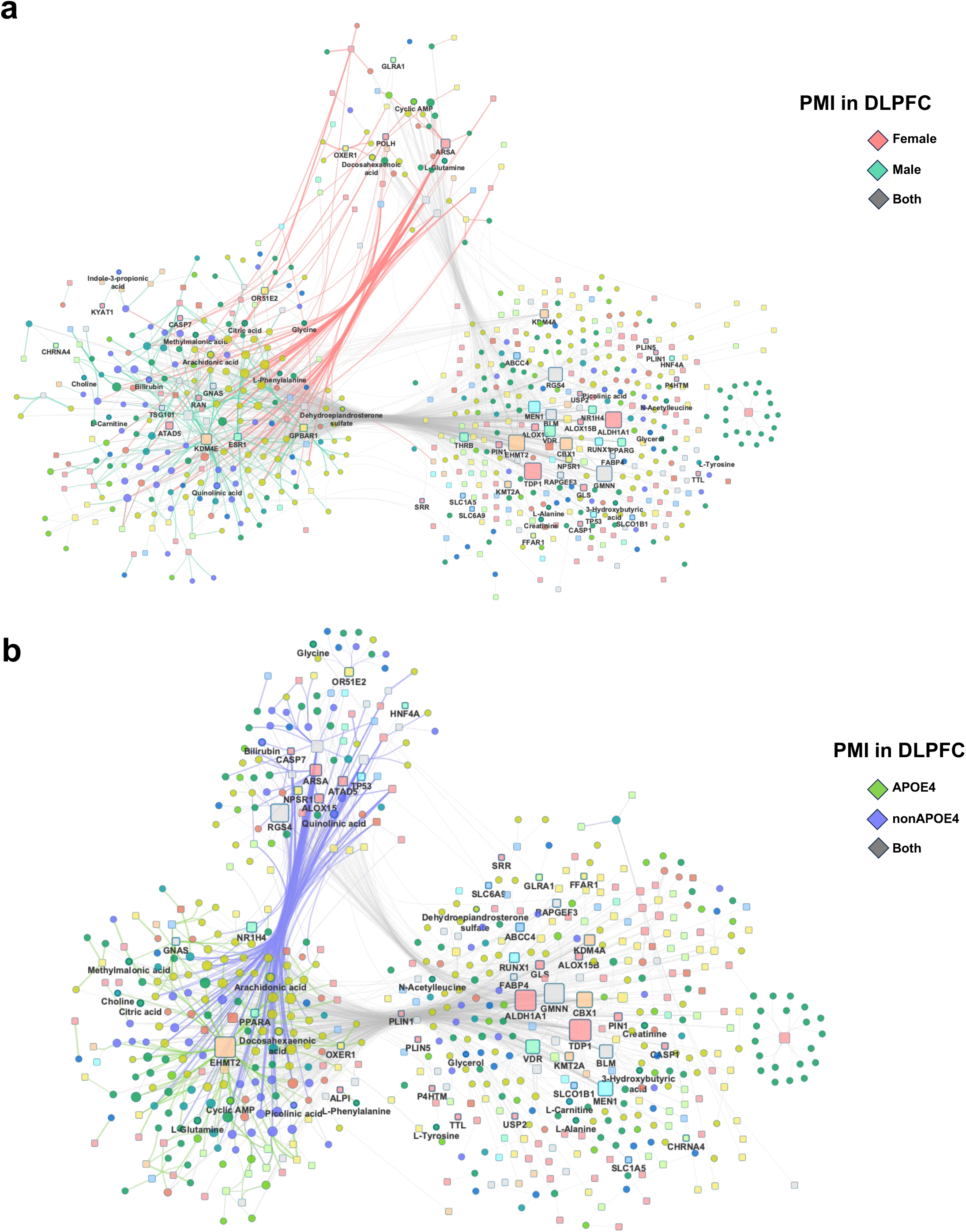
Genetic-supported metabolites-mediated metabolite-sensor network by AD risk factors in the DLPFC. **a,b** Network depicting sex-specific metabolite-sensor pairs (**a**) and APOE4-specific pairs (**b**). Line colors indicate the sex-specific or APOE4-specific pairs. Metabolites are shown in circle and proteins are shown in square. MR-prioritized metabolites and their pairs were labeled. The colors or labels styles are consistent with Figure 6.

**Figure S11.**
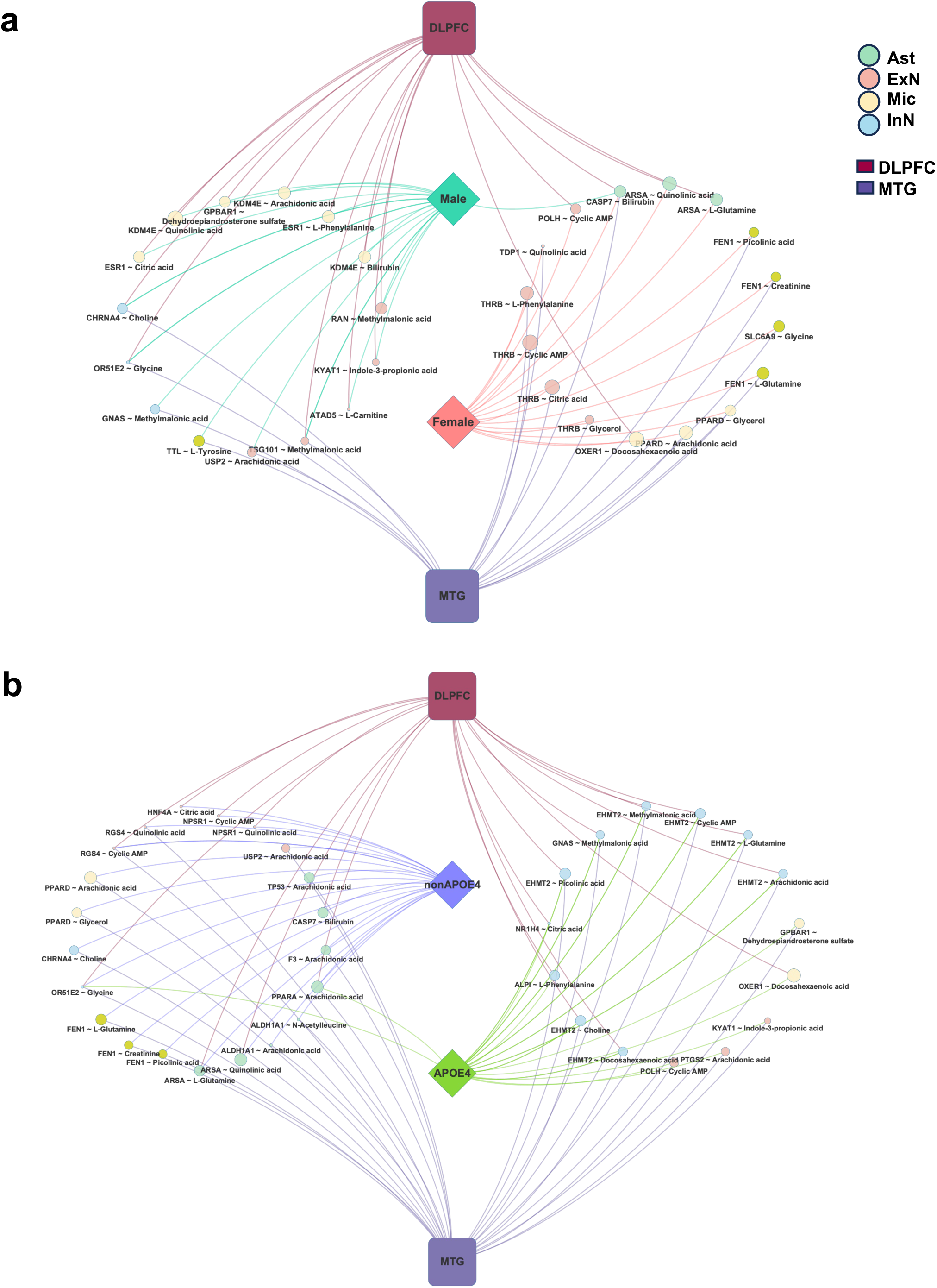
Differences of genetic-supported metabolites-mediated metabolite-sensor network across AD risk factors. **a,b** Network representations of sex-specific (**a**) or APOE4-specific (**b**) AD likely causal pairs in both MTG and DLPFC regions.

